# Neuroanatomical dissection of the MC3R circuitry regulating energy rheostasis

**DOI:** 10.1101/2024.04.22.590573

**Authors:** Ingrid Camila Possa-Paranhos, Jared Butts, Emma Pyszka, Christina Nelson, Dajin Cho, Patrick Sweeney

## Abstract

Although mammals resist both acute weight loss and weight gain, the neural circuitry mediating bi-directional defense against weight change is incompletely understood. Global constitutive deletion of the melanocortin-3-receptor (MC3R) impairs the behavioral response to both anorexic and orexigenic stimuli, with MC3R knockout mice demonstrating increased weight gain following anabolic challenges and increased weight loss following anorexic challenges (i.e. impaired energy rheostasis). However, the brain regions mediating this phenotype remain incompletely understood.

Here, we utilized MC3R floxed mice and viral injections of Cre-recombinase to selectively delete MC3R from medial hypothalamus (MH) in adult mice. Behavioral assays were performed on these animals to test the role of MC3R in MH in the acute response to orexigenic and anorexic challenges. Complementary chemogenetic approaches were used in MC3R-Cre mice to localize and characterize the specific medial hypothalamic brain regions mediating the role of MC3R in energy homeostasis. Finally, we performed RNAscope in situ hybridization to map changes in the mRNA expression of MC3R, POMC, and AgRP following energy rheostatic challenges.

Our results demonstrate that MC3R deletion in MH increased feeding and weight gain following acute high fat diet feeding in males, and enhanced the anorexic effects of semaglutide, in a sexually dimorphic manner. Additionally, activation of DMH MC3R neurons increased energy expenditure and locomotion. Together, these results demonstrate that MC3R mediated effects on energy rheostasis result from the loss of MC3R signaling in the medial hypothalamus of adult animals and suggest an important role for DMH MC3R signaling in energy rheostasis.

**Key Points:**

- MC3R signaling regulates energy rheostasis in adult mice
- Medial hypothalamus regulates energy rheostasis in adult mice
- Energy rheostasis alters mRNA levels of AgRP and MC3R in DMH
- DMH MC3R neurons increase locomotion and energy expenditure
- MC3R expression in DMH is sexually dimorphic

## 1. Introduction

In mammals body weight is remarkably stable in the short term (i.e. over the course of days), as humans and rodents will actively resist acute weight gain or weight loss. In response to energy deprivation, changes in feeding, energy expenditure, and neuroendocrine circuits occur to resist against severe weight loss(1–8). While adaptive in preventing starvation, these changes contribute to the difficulty in maintaining weight loss(8,9). Conversely, both humans and rodents defend against acute weight gain by engaging behavioral, autonomic, and neuroendocrine circuitry to prevent excessive weight gain(1,10,11). For example, in human’s periods of excessive weight gain, such as holiday-induced weight gain, are typically followed by weight loss and a return to prior body weight(12). Consistently, forced overfeeding in rodents and primates (over 1-2 weeks) via intragastric infusions is followed by drastic anorexia until body weight returns to the levels observed prior to overfeeding(1,11). Although the mechanisms preventing excessive weight loss are relatively well understood, the mechanisms preventing excessive weight gain are largely unknown. Further, since excessive weight gain or weight loss are typically studied in isolation, it remains unclear how neural circuits adapt in animals to resist both orexigenic and anorexic challenges.

Based on prior reports, the melanocortin-3 receptor (MC3R) exerts a unique role in energy homeostasis, with MC3R knockout mice exhibiting an impaired response to both acute weight loss and weight gain, which was termed altered energy rheostasis(13–15). MC3R knockout (KO) mice consume normal amounts of food on a regular chow diet when fed *ad libitum*, but gain excessive weight upon access to a high fat diet or surgical ovariectomy(14). Conversely, these mice are hypersensitive to various anorexic challenges, including stress-related anorexia(16) (restraint stress and social-isolation induced anorexia), diverse forms of pharmacological anorexia(16,17), and physiological anorexia(15,18)(tumor-associated anorexia). Together, these findings suggest that MC3R controls “energy rheostasis”, or the magnitude of metabolic responses in both the positive and negative direction following anabolic or catabolic stimuli(14). Thus, in contrast to other mutations in the leptin-melanocortin pathway, which result in uncontrolled hyperphagia and weight gain regardless of the dietary condition, deletion of MC3R results in an exaggerated response in opposing directions to orexigenic or anorexic stimuli.

The leptin-melanocortin pathway is among the most well studied circuitry regulating feeding and body weight. This pathway is essential for regulating feeding and energy expenditure and mutations in genes related to the leptin-melanocortin circuit are the most common cause of monogenic obesity in humans(19). The melanocortin system is composed of pro-opiomelanocortin (POMC) and agouti-related peptide (AgRP) neurons in the arcuate nucleus of the hypothalamus, which produce the endogenous agonist (aMSH) and antagonist (AgRP) for the central melanocortin receptors (MC3R and MC4R)(19–21). AgRP neurons are activated by caloric deprivation (22–24), resulting in the release of the melanocortin receptor antagonist/inverse agonist AgRP (in addition to the inhibitory neurotransmitter GABA and neuropeptide Y)(19,25) to stimulate feeding and reduce energy expenditure. Conversely, POMC neurons are activated by caloric sufficiency (26) and release α-melanocyte stimulating hormone (α-MSH), an agonist at the melanocortin-4 receptor (MC4R) and melanocortin-3 receptor (MC3R)(19) to suppress feeding and increase energy expenditure.

AgRP and POMC neurons mediate their effects on feeding and energy expenditure by projecting to multiple downstream brain regions(19,21), where they act on secondary neurons containing MC3R and/or MC4R (in addition to non-MC3R and non-MC4R expressing neurons via GABA and/or NPY). MC4R is thought to be the primary receptor mediating satiety downstream of POMC neurons(27), while the role of MC3R is energy homeostasis is less well understood(28,29). Consistent with this interpretation, rodents and humans with mutations in *MC4R* are hyperphagic and obese(30–33), while *MC3R* knockout mice exhibit minor late onset obesity(28,29), without noticeable hyperphagia when provided *ad libitum* access to regular chow diet. Recently a case of homozygous loss-of function of MC3R was reported in humans, with this patient displaying obesity, increased fat mass, reduced lean mass, and a delayed puberty phenotype like those previously characterized in MC3R KO mice(34), suggesting that the function of MC3R may be conserved across species.

Although prior work implicates MC3R in excessive responses to both orexigenic and anorexic stimuli (energy rheostasis), this interpretation is almost entirely based on behavioral data in which the MC3R has been deleted globally from development. Constitutive deletion of MC3R results in a multitude of secondary metabolic and neuroendocrine abnormalities, such as increased fat mass and leptin levels, and impaired levels of corticosterone and thyroid hormone(13,14,28,29), which confound the interpretation of behavioral phenotypes in MC3R KO mice since these changes may alter behavioral phenotypes independent of MC3R action. Thus, it remains unclear if central MC3R signaling regulates energy rheostasis or if the previously reported energy rheostasis phenotype is a result of secondary abnormalities observed in constitutive MC3R KO mice. Furthermore, prior studies indicate that global gene deletions can result in developmental compensation, such that a limited behavioral phenotype is observed following constitutive deletion. For example, despite a well-established critical role in energy homeostasis, constitutive deletion of important metabolic receptors such as glucagon-like-1 receptor(35), ghrelin(36), and ghrelin receptor(37) result in limited metabolic phenotypes. Therefore, the relative importance of central MC3R signaling in adult animals cannot be ascertained solely from global KO mouse models.

Recent work demonstrates that MC3R is widely expressed throughout the brain, with dense MC3R expression observed throughout all major brain regions. Of interest to the role of MC3R in energy rheostasis, particularly strong MC3R expression is observed in medial hypothalamic structures (arcuate nucleus, ventral-medial hypothalamus, dorsal-medial hypothalamus)(38). However, given the dense brain-wide expression of MC3R, the specific brain regions and neural circuits mediating the role of MC3R in energy rheostasis are largely unknown. In the present study we utilized a viral deletion approach to specifically delete the MC3R within select MH regions in adult mice. Our findings indicate that adult specific deletion of MC3R throughout the MH recapitulates many of the energy rheostasis phenotype observed in MC3R KO mice. Surprisingly, despite a critical role for MC3R in the arcuate nucleus in feeding(13,14,16,17), we find that the ability of MC3R to control energy rheostasis does not require the arcuate nucleus. Further, we identify the dorsal-medial hypothalamus (DMH) as an important site for MC3R-mediated changes in energy rheostasis and characterize a specific role for DMH MC3R neurons in controlling energy expenditure and locomotion.

## 2. Results

### 2.1. Deletion of MC3R in the medial hypothalamus (MH) does not alter regular chow feeding

Although MC3R expression is observed throughout the brain, particularly strong expression is observed in medial hypothalamic (MH) regions (including the arcuate nucleus, ventral-medial hypothalamus, and dorsal medial hypothalamus) (38). Given the established role of MH regions in energy homeostasis, we first sought to determine the role of MH MC3R signaling in energy homeostasis in adult mice. To selectively delete MC3R in the MH of male and female mice, we injected adeno-associated virus (AAV) expressing Cre recombinase into the MH in mice containing loxp sites flanking the entire exon encoding MC3R (MC3R floxed mice) (**Figures 1A, 1B, and S1**). To control for any potential off-target effects of Cre expression, WT littermate mice were also injected with the identical Cre expressing virus. First, to validate successful MC3R deletion, we performed RNAscope in situ hybridization analysis to quantify MC3R mRNA expression in the MH in MC3R floxed and WT mice injected with AAV-Cre virus. Consistent with prior reports(38), we identified dense MC3R mRNA expression throughout the MH in WT mice injected with AAV-Cre (**Figure 1C**). In contrast, AAV-Cre injections in MC3R floxed mice markedly reduced MC3R mRNA expression throughout the MH, indicating successful viral-mediated deletion of MC3R (**Figures 1D and 1E**).

**Figure 1:**
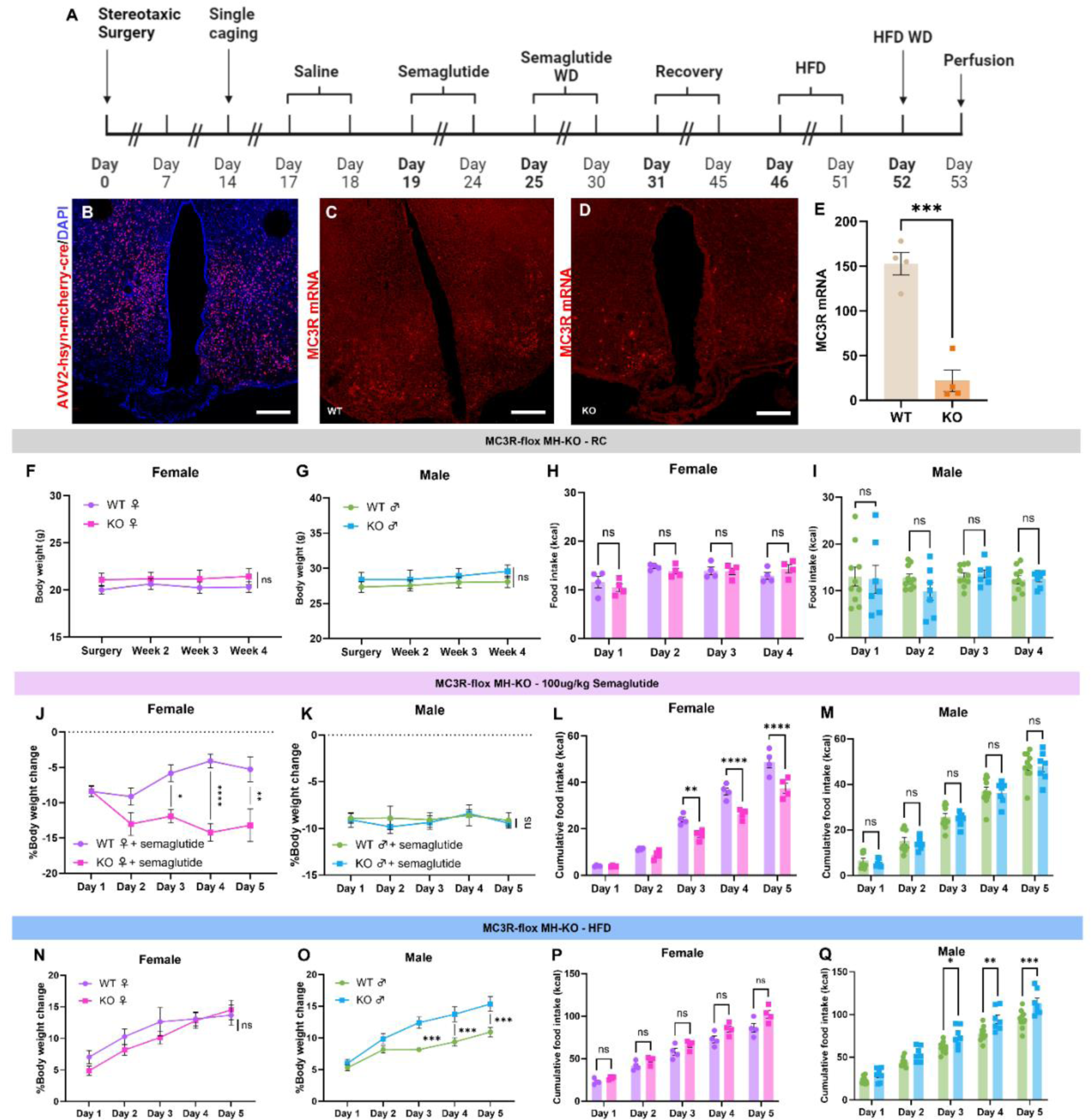
Adult hypothalamic deletion of MC3R alters energy rheostasis. (A) Timeline of experimental protocol for experiments in Figure 1. Created with Biorender. (B) Viral location example of a MC3R-flox mouse with AVV2-hsyn-mcherry-cre in medial hypothalamus (MH). Scale bars in 300um. (C) RNAscope image of example WT mouse injected with AVV2-hsyn-mcherry-cre and with MC3R mRNA in red. Scale bars in 300um. (D) RNAscope image of representative MC3R-flox mouse injected with AVV2-hsyn-mcherry-cre and with MC3R mRNA shown in red. Sections shown in 1B and 1D come from the same mouse and adjacent sections. Note the increased MC3R mRNA expression on the bottom left in 1D corresponds well with the limited AAV-cre viral spread in this region shown in 1B. Scale bars in 300um. (E) MC3R mRNA count from RNAscope sections in WT and MH MC3R KO mice (unpaired t-test, p***= 0.0003, n=4 mice/group). (F and G) WT and MC3R-MH KO body weight curves from surgery until 4 weeks post-surgery. (F) Female mice (G) Male mice (F: two-way ANOVA with multiple comparisons, p=0.6774 n=4 mice/group G: two-way ANOVA with multiple comparisons p=0.7019 n=10 mice for WT group and 7 mice for MH MC3R floxed group). (H and I) Food intake for 4 days, 2 weeks after surgery (H) Female mice (I) Male mice (H: two-way ANOVA with multiple comparisons p=0.5798 n=4 mice/group; I: two-way ANOVA with multiple comparisons, p>0.9999 n=10 mice for WT group and 7 mice for MH MC3R floxed group). (J and K) Daily percentage body weight change compared to saline day after daily 100mg/kg injections of semaglutide (J) Female mice (K) Male mice (J: baseline-corrected followed by two-way ANOVA with multiple comparisons, p*=0.0164, p**=0.0012, p****=<0.0001 n=4 mice/group; K: baseline-corrected followed two-way ANOVA with multiple comparisons, p=0.9991 n=10 mice in WT group and 7 mice in MH KO group). (L and M) 24-hour cumulative daily food intake during semaglutide administration days (L) Female mice (M) Male mice (L: two-way ANOVA with multiple comparisons, p**<0.0074, p****=<0.0001 n=4 mice/group; M: two-way ANOVA with multiple comparisons, p>0.9999 n=10 mice for WT group and 7 mice for MH KO group). (N and O) Percentage body weight curve, comparing WT to MC3R-flox MH-KO with *ad libitum* access to HFD (N) Female mice (O) Male mice (N: two-way ANOVA with multiple comparisons p=0.9985 n=4 mice/group; O: two-way ANOVA with multiple comparisons, p***<0.0007 n=10 mice for WT group and 7 for MH MC3R KO group). (P and Q) Cumulative HFD food intake in WT and MC3R-flox MH-KO mice (P) Female mice (Q) Male mice (P: two-way ANOVA with multiple comparisons p=0.1385 n=4 mice/group; Q: two-way ANOVA with multiple comparisons, p*=0.0151, p**=0.0025, p***=0.0004 n=10 mice in WT group and 7 mice in MH KO group). Data points represent individual mice.

Given the previous literature implicating MC3R in energy rheostasis(14), we designed an experimental protocol to assess the role of MC3R signaling in MH in mediating acute responses to anorexic and orexigenic stimuli in adult mice (**Figure 1A**). First, we measured feeding behavior and body weight change in MH MC3R KO and WT mice in animals provided *ad libitum* access to a regular chow diet (**Figure 1A**). Deletion of MC3R in the MH did not alter body weight in male or female mice (**Figures 1F and 1G**) and had no effect on food intake in mice provided *ad libitum* access to a regular chow diet (**Figures 1H and 1I**). Thus, like global developmental deletion of MC3R, adult medial hypothalamic deletion of MC3R does not affect feeding or body weight in basal conditions(14).

2.2. **MH deletion of MC3R increases anorexic response to semaglutide.**

MC3R KO mice are hypersensitive to diverse anorexic stimuli including stress-related anorexia (restraint stress and social isolation)(16), pharmacological anorexia(14,16,17) (i.e. administration of GLP1R agonists and other anorexic compounds), and physiological anorexia(15,18) (i.e. tumor cachexia). Therefore, following daily habituation to saline injections, we next tested if MH deletion of MC3R alters the anorexic response to the GLP1R agonist semaglutide(39,40), as observed in global MC3R KO mice(14,17). Female MC3R MH-KO mice lost significantly more weight than WT littermates (**Figure 1J**) and ate significantly less during semaglutide treatment (**Figure 1L**). In contrast, no difference in body weight or feeding were observed between male WT mice and male MH MC3R KO mice during semaglutide administration (**Figures 1K and 1M**). Following semaglutide treatment, female MC3R MH-KO mice maintained a significantly lower body weight for 3 days after the withdrawal (**Figure S2A**), with no significant difference in their food intake (**Figure S2C**), while no phenotype was observed in males following semaglutide administration (**Figures S2B and S2D**).

### 2.3. MH deletion of MC3R increases body weight gain on a high fat diet

Prior studies indicate that MC3R KO mice gain more weight on a high fat diet(14). Therefore, following washout and recovery from semaglutide treatment (**Figure 1A**), we next tested if MH KO of MC3R increases feeding and body weight when mice are provided acute access to a palatable high-fat diet (HFD). During *ad libitum* access to HFD, male MH KO mice gained significantly more weight starting on the third day of access to HFD until the end of the experiment (**Figure 1O**), which was accompanied by increased HFD food intake (**Figure 1Q**). In contrast, female MH KO mice did not differ from WT littermates in both body weight gain (**Figure 1N**) or food intake (**Figure 1P**). Thus, MH deletion of MC3R increases body weight gain and food intake in male mice fed a high fat diet, recapitulating the phenotype previously observed in male mice with global deletion of MC3R(14).

Recent reports demonstrate that acute access to HFD rapidly alters the activity of hypothalamic AgRP neurons, resulting in a de-valuation of regular chow diet, such that acute anorexia develops when animals are switched back from a HFD to a regular chow diet(41). Since AgRP neurons produce the endogenous antagonist for the MC3R (AgRP), and prior work indicates that MC3R KO mice are hypersensitive to anorexic stimuli, we next measured the feeding and weight loss response in WT and MH MC3R KO mice after switching the diet back to a regular chow diet. Both female WT and MH MC3R KO mice exhibited a similar anorexic response upon switching to a regular chow diet, losing a similar amount of weight following the switch to regular chow (**Figure S2E**) and equivalently reducing the kilocalories ingested after the transition to regular chow diet (**Figure S2G**). In contrast, male MH KO mice lost less body weight (**Figure S2F**) and had a reduced anorexic response compared to WT mice following the transition to a regular chow diet (**Figure S2H**).

### 2.4 Energy rheostatic challenges alter mRNA expression of AgRP and MC3R in MH

MC3R signaling is bi-directionally regulated by the endogenous melanocortin receptor agonist aMSH (produced by the POMC peptide in POMC neurons) which stimulates MC3R, and the endogenous MC3R antagonist AgRP, which inhibits the MC3R. Given the role of MC3R signaling in energy rheostasis, we next utilized RNAscope in situ hybridization to quantify the mRNA expression of AgRP, POMC, and MC3R in the MH in the context of the energy rheostatic assays previously described (acute HFD administration and acute semaglutide administration). WT male mice were either provided *ad libitum* access to regular chow diet, HFD for five days, or 5 days of semaglutide treatment, and processed for in situ hybridization analysis of AgRP, POMC, and MC3R in MH. Since fasting increases mRNA expression of AgRP(24,42), we also quantified AgRP and POMC expression in a separate cohort of mice that were fasted for 16 hours to validate the sensitivity of RNAscope analysis for detecting transcriptional changes in AgRP and POMC. As expected, fasting significantly increased mRNA expression of AgRP, and non-significantly reduced POMC expression (**Figure S3A-S3F**). No significant difference in POMC expression was observed following semaglutide treatment or HFD administration (**Figures 2A-2D**). Although semaglutide administration did not alter mRNA levels of AgRP, acute HFD feeding significantly reduced AgRP expression (**Figures 2E-2H**), suggesting that reduced AgRP levels may contribute to the role of MC3R in regulating the acute orexigenic response to high fat diet.

**Figure 2:**
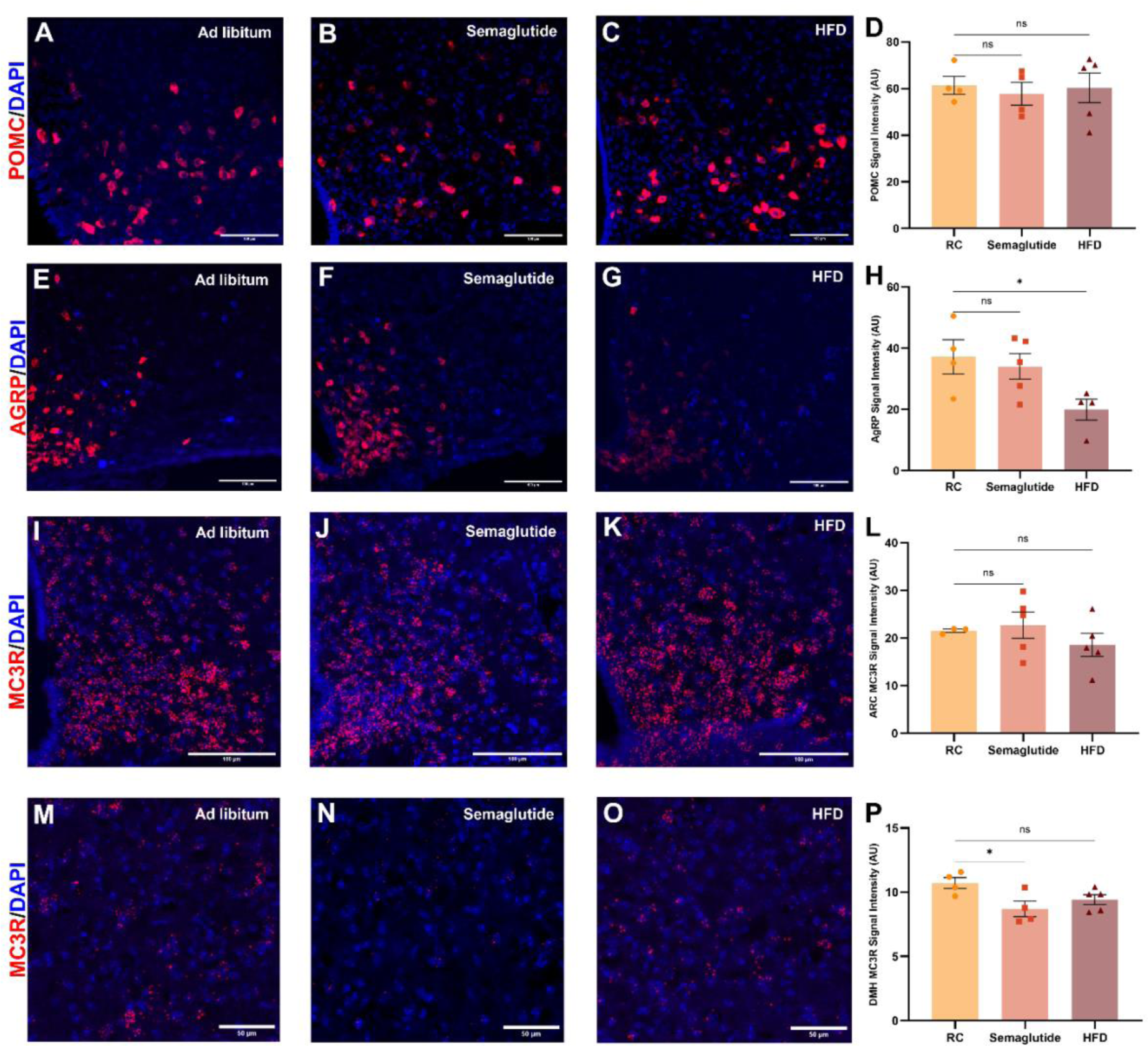
Semaglutide alters expression of MC3R mRNA in DMH. (A, B, and C) Confocal image of POMC mRNA expression (red) with DAPI (blue) of a mouse (A) with a*d libitum* access to regular chow (RC), (B) after 5 days of semaglutide administration and (C) after 5 days of high fat diet (HFD) access. Scale bars 100um. (D) Comparison between the intensity of POMC mRNA signal during a*d libitum*, semaglutide administration for 5 days, and access to HFD for 5 days. (two-way ANOVA with multiple comparisons, RC-Semaglutide p=0.8615; RC-HFD p=0.9870). (E, F, and G) Confocal image of AgRP mRNA expression (red) with DAPI (blue) of a mouse (E) with *ad libitum* access to RC, (F) after 5 days of semaglutide administration, and (G) after 5 days of HFD access. Scale bar 100um. (H) Comparison of the intensity of AgRP mRNA signal during *ad libitum*, semaglutide administration for 5 days, and access to HFD for 5 days. (two-way ANOVA with multiple comparisons, RC-Semaglutide p=0.8322; RC-HFD p*=0.0452). (I, J, and K) Confocal image of MC3R mRNA (red) expression in ARC with DAPI (blue) of a mouse (I) with *ad libitum* access to RC, (J) after 5 days of semaglutide administration (K) after 5 days of HFD access. Scale bar 100um. (L) Comparison of the intensity of MC3R mRNA signal in ARC during *ad libitum*, semaglutide administration for 5 days, and access to HFD for 5 days. (two-way ANOVA with multiple comparisons, RC-Semaglutide p=0.9273; RC-HFD p=0.6368). (M, N, and O) Confocal image of MC3R mRNA (red) expression in DMH with DAPI (blue) of a mouse with (M) *ad libitum* access, (N) after 5 days of semaglutide administration (O) after 5 days of HFD access. Scale bar=50um. (P) Comparison of the intensity of MC3R mRNA signal in DMH during *ad libitum*, semaglutide administration for 5 days and access to HFD for 5 days. (two-way ANOVA with multiple comparisons, RC-Semaglutide p*=0.0277; RC-HFD p=0.1343). Data points represents the average of the signal intensity from all the sections of each individual animal.

Since MC3R signaling regulates the acute anorexic and orexigenic response to semaglutide and HFD, we next quantified if the mRNA expression of MC3R was altered within the MH (arcuate nucleus, ventral medial hypothalamus, and dorsal-medial hypothalamus) following acute semaglutide or HFD administration. Both HFD and semaglutide administration had no effect on mRNA expression of MC3R within the arcuate nucleus (**Figures 2I-2L**) and VMH (**Figures S3G-S3J**). However, semaglutide administration reduced MC3R mRNA expression in the DMH, while HFD feeding did not alter MC3R expression in DMH (**Figures 2M-2P**).

### 2.5 Role of MH MC3R signaling in energy rheostasis does not require arcuate nucleus

The MH is comprised of multiple sub-regions (arcuate nucleus, ventral-medial hypothalamus, and dorsal-medial hypothalamus) that each have critical roles in energy homeostasis(43). Importantly, MC3R is expressed throughout the entire MH(38), with especially high expression observed in the arcuate nucleus(38). Most prior work indicates that the primary site of action for MC3R-mediated effects on feeding behavior is the arcuate nucleus of the hypothalamus, where MC3R acts to both control the activity level of AgRP neurons and regulate presynaptic release from AgRP neuron terminals(14,44). However, MC3R expression is also observed in the ventral-medial hypothalamus (VMH)(38,45,46) and dorsal-medial hypothalamus (DMH)(38), brain regions that are critically involved in feeding and energy homeostasis and where the function of MC3R is less well understood. Further, MC3R mRNA expression is reduced in DMH following semaglutide treatment (**Figures 2N and 2P**), suggesting that DMH may also contribute to the role of MC3R in energy rheostasis. We therefore next examined if MC3R signaling in the dorsal portions of medial hypothalamus (dMH) contribute to energy rheostasis. In new cohorts of mice, AAV-Cre virus was targeted more dorsally in the MH in WT and MC3R floxed mice to determine if the role of MH MC3R signaling in energy rheostasis requires the arcuate nucleus (**Figures 3A, 3B and S4**). Viral expression was primarily localized to the dorsal-medial hypothalamic area, with more sparse expression also observed in the ventral medial hypothalamus and posterior hypothalamus in some mice (subsequently referred to as dMH deletion of MC3R, **Figures 3B and S4**). However, no viral expression was observed in the arcuate nucleus or lateral hypothalamus (**Figure S4**). As previously described (**Figure 3A**) we next characterized the response of dMH-MC3R KO and WT mice to acute anorexic and orexigenic challenges (**Figures 3 and 4**). As observed with MH deletion of MC3R, deletion of MC3R in the dMH did not alter body weight or food intake in male or female mice on a regular chow *ad libitum* diet (**Figures 3C-3F**). Since MC3R KO mice have a defective orexigenic response to both fasting and caloric restriction(14), we first tested if MC3R signaling in dMH also regulates weight regain following caloric restriction. During 70% caloric restriction both WT and dMH MC3R KO mice lost similar amounts of weight (**Figures 3G and 3H**). Following re-feeding in WT and dMH MC3R KO mice, both groups regained weight at a similar pace (**Figures 3G and 3H**). Thus, in contrast to MC3R signaling in arcuate AgRP neurons, dMH MC3R signaling does not regulate weight regain following caloric restriction(14).

**Figure 3:**
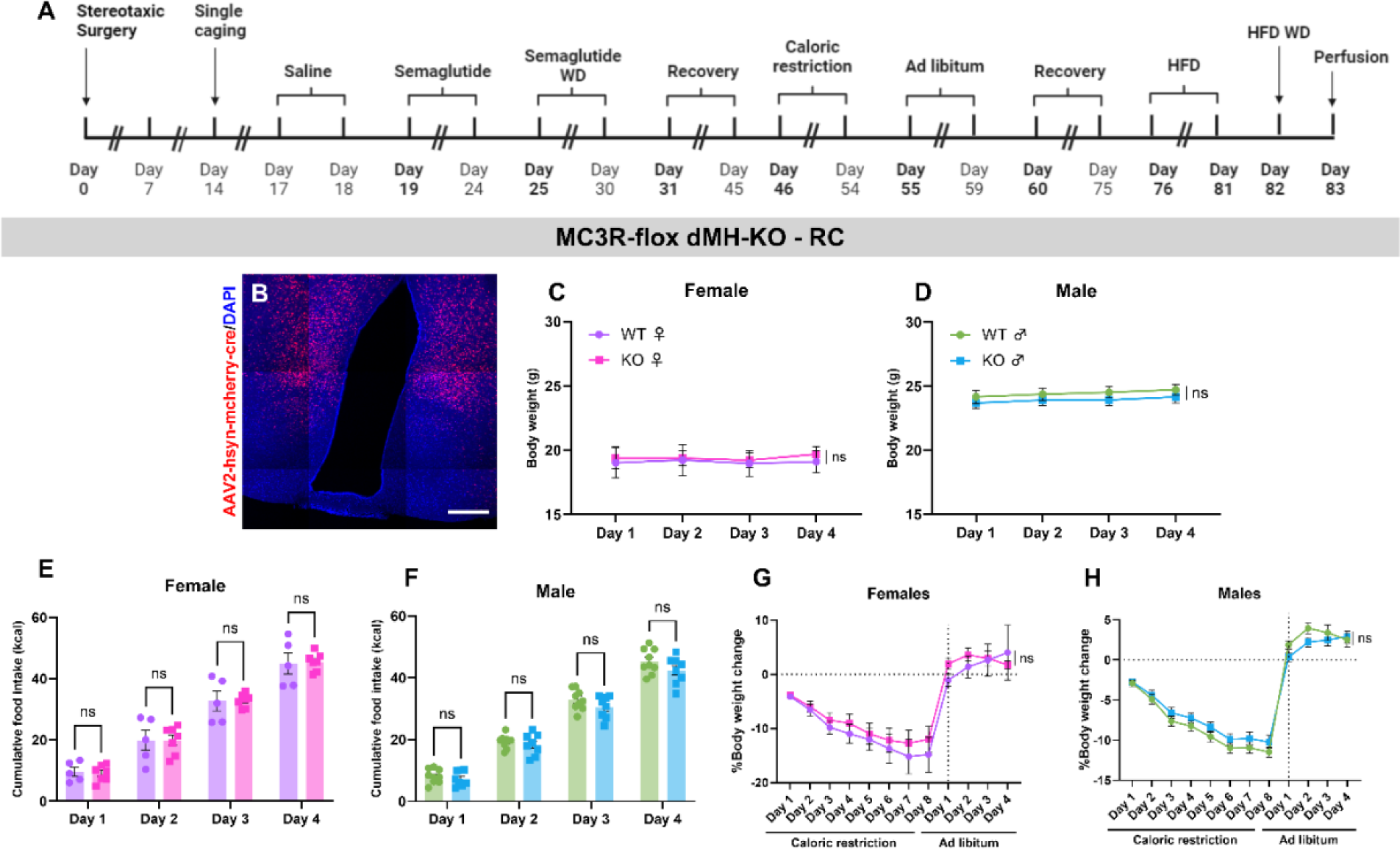
Deletion of MC3R in DMH does not alter the food intake or body weight on regular chow diet. (A) Timeline of experimental protocol for experiments in Figures 3 and 4. Created with Biorender. (B) Example image of a MC3R-flox mouse with AVV2-hsyn-mcherry-cre in dMH. Scale bar 300um. (C and D) Body weight curve with *ad libitum* access to regular chow 2 weeks after surgery (C) Female mice (D) Male mice (C: two-way ANOVA with multiple comparisons, p=0.9723 n=5 mice for WT group and 7 mice for dMH KO group; D: two-way ANOVA with multiple comparisons, p=0.8486 n=9 mice for WT group and 8 mice for dMH KO group). (E and F) Cumulative regular chow food intake 2 weeks after surgery (E) Female mice (F) Female mice (E: two-way ANOVA with multiple comparisons, p>0.9999 n=5 mice for WT group and 7 mice for dMH KO group; F: two-way ANOVA with multiple comparisons, p=0.5435 n=9 mice for WT group and 8 mice for dMH KO group). (G and H) Percentage body weight change during caloric restriction, using the body weight from last day of *ad libitum* access to regular chow as the baseline (G) Female mice (H) Male mice (G: baseline-corrected followed by two-way ANOVA with multiple comparisons, p>0.9999 n=5 mice for WT group and 7 mice for dMH KO group, H: baseline-corrected followed by two-way ANOVA with multiple comparisons, p>0.9999 n=9 mice for WT group and 8 mice for dMH KO group). Data points represent individual mice.

**Figure 4:**
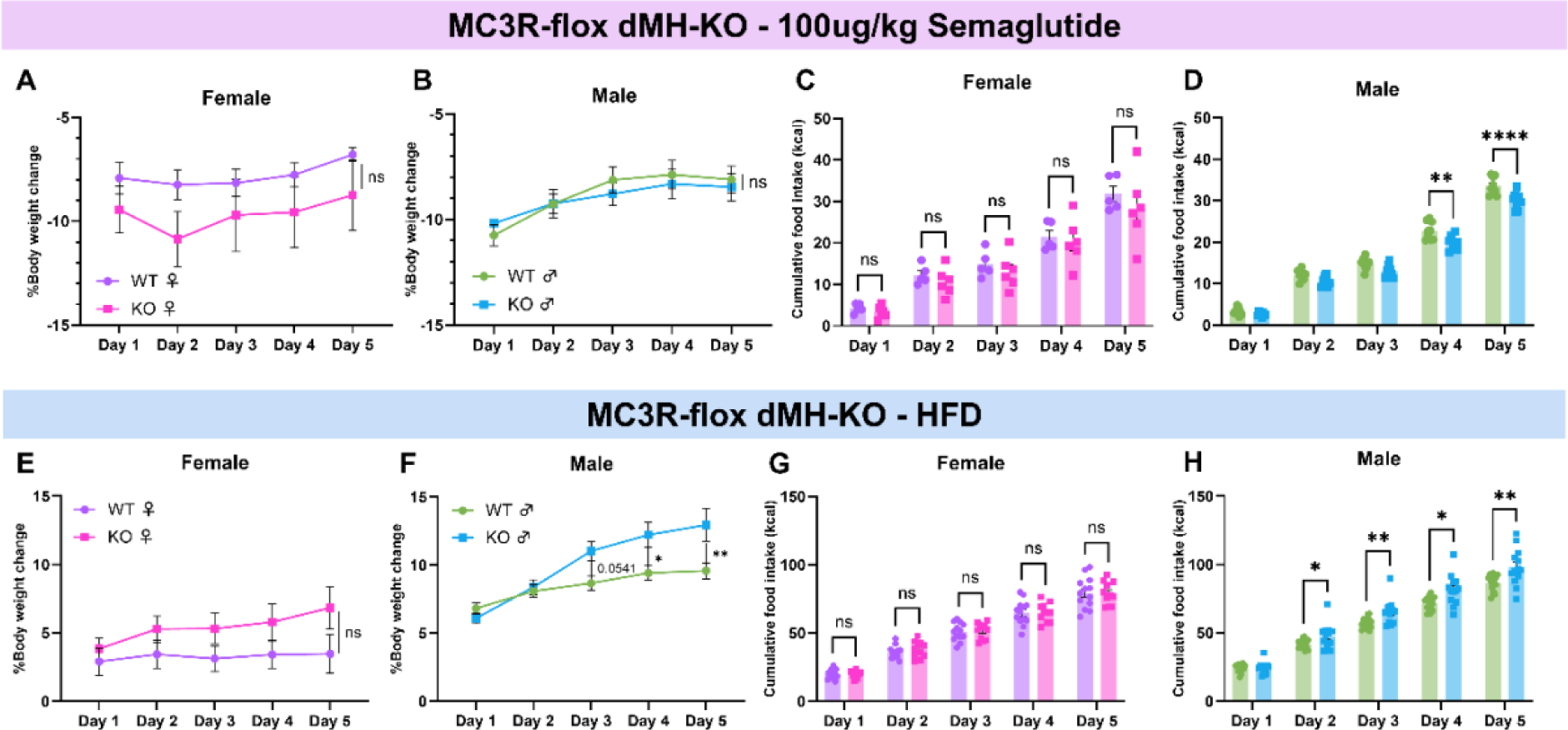
dMH MC3R signaling regulates energy rheostasis. (A and B) Daily percentage body weight change using saline as the baseline after 100mg/kg of semaglutide administration (A) Female mice (B) Male mice (A: baseline-corrected followed by two-way ANOVA with multiple comparisons p=0.8071 n=5 mice in WT group and 6 mice in dMH KO group; B: baseline-corrected followed by two-way ANOVA with multiple comparisons, p=0.9958 n=9 WT mice and 8 dMH KO mice). (C and D) 24-hour cumulative food intake after daily semaglutide administration (C: two-way ANOVA with multiple comparisons, p=0.9249 n=5 mice in WT group and 6 mice in dMH KO group; D: two-way ANOVA with multiple comparisons, p**=0.0011, p****<0.0001 n=9 WT mice and 8 dMH KO mice). Data points represent individual mice. (E and F) Percentage of body weight change during access to *ad libitum* HFD, using the last day of regular chow as the baseline (E) Female mice (F) Male mice (E: baseline-corrected followed by two-way ANOVA with multiple comparisons, p=0.174 n=13 WT mice and 11 dMH KO mice; F: baseline-corrected followed by two-way ANOVA with multiple comparisons, p*=0.0139, p**=0.0017 n=16 WT mice and 13 dMH KO mice) (G and H) Cumulative daily food intake during HFD access (G) Female mice (H) Male mice (G: statistical analysis: two-way ANOVA with multiple comparisons, p=0.9691 n=13 WT mice and n=11 dMH KO mice; H: two-way ANOVA with multiple comparisons, p*<0.0473, p**<0.0094 n=16 WT mice and 13 dMH KO mice).

Following recovery from caloric restriction, we next tested if dMH deletion of MC3R alters the anorexic response to anorexigenic stimuli by administering semaglutide or vehicle to WT and dMH MC3R KO mice. No difference in body weight was detected between female WT and dMH MC3R KO mice following daily semaglutide administration (**Figures 4A and 4C**). In contrast, male MC3R dMH-KO mice consumed significantly less food than WT littermates during semaglutide treatment (**Figure 4D**), although no difference in BW was detected (**Figure 4B**). No difference was observed in the rate of body weight re-gain or feeding between WT and dMH MC3R KO mice following semaglutide withdrawal (**Figures S5A-S5D**).

Following recovery from semaglutide treatment (**Figure 3A**), we next tested if dMH deletion of MC3R alters feeding and body weight when mice are provided a high fat diet. Male MC3R dMH KO mice had a significant increase in body weight and food intake following HFD administration, relative to WT littermate mice (**Figures 4F and 4H**). However, this effect was not observed in female mice, although female MC3R dMH-KO mice demonstrated a tendency towards increased body weight gain on HFD (**Figures 4E and 4G**). To determine if dMH MC3R deletion alters the anorexic response associated with switching HFD-fed mice to a regular chow diet, we measured feeding and body weight after changing HFD fed mice to a regular chow diet. Both dMH MC3R KO and WT littermate mice demonstrated a similar anorexic and weight loss response to withdrawal from HFD (**Figures S5E-S5H**). Thus, although the arcuate nucleus exerts an important role in energy rheostasis, additional regions outside the arcuate nucleus (i.e. dMH area) are also capable of regulating energy rheostasis.

### 2.6 dMH MC3R signaling selectively regulates the acquisition of weight gain on HFD

Our previous results demonstrate that dMH deletion of MC3R increases weight gain in the first week of high fat diet administration (**Figures 4F and 4H**). However, it is unclear if deletion of MC3R specifically regulates the initial acquisition of weight gain on HFD, or if deletion of MC3R amplifies weight gain in mice that already exhibit diet-induced obesity. To test this, in new cohorts of mice, we fed WT littermate and MC3R floxed mice with a HFD for six weeks to induce obesity (**Figure 5A**). Importantly, no difference in initial weight gain or feeding was detected between WT and MC3R floxed mice prior to AAV injections, indicating that the presence of the floxed allele does not alter the baseline response to high fat diet (**Figures 5B and 5C**). Further, both groups of mice developed obesity at a similar rate (**Figures 5B and 5C**). After six weeks of HFD, we injected AAV-Cre virus into the dMH area in WT and MC3R floxed mice and continued to measure feeding and body weight gain for three additional weeks. Like prior dMH targeted viral injections, AAV expression was primarily localized to the dorsal-medial hypothalamic regions, with sparse expression observed in the nearby posterior hypothalamus (**Figure S6**). In contrast to the acute response to initial HFD administration, dMH deletion of MC3R does not alter feeding or body weight gain when mice were accustomed to HFD and had already developed diet-induced obesity (**Figures 5B-5E**). Thus, MC3R signaling in dMH regulates the initial acquisition of weight gain in response to high fat diet but does not amplify weight gain following diet-induced obesity.

**Figure 5.**
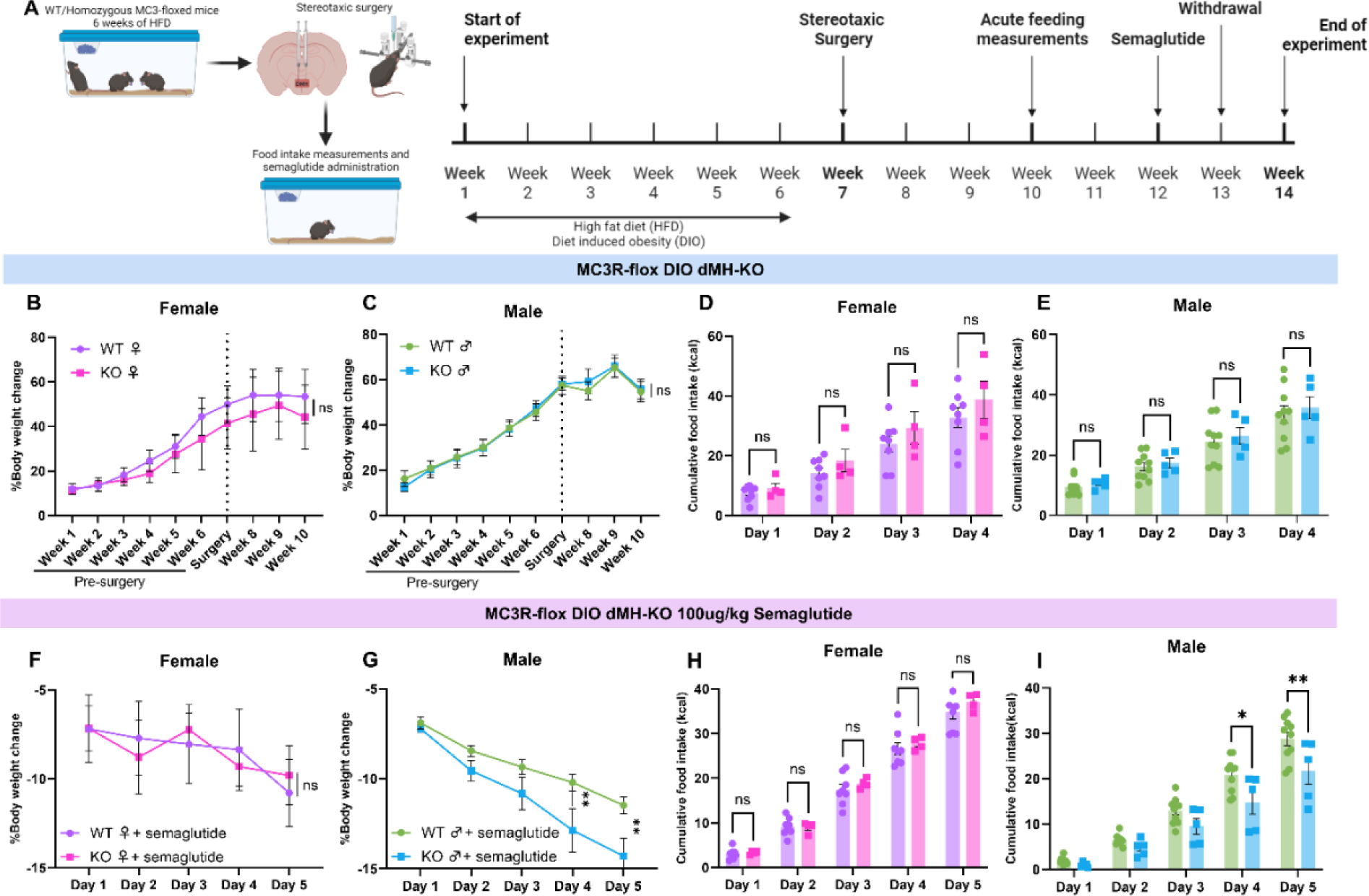
Adult DMH deletion of MC3R selectively regulates the acquisition of body weight on high fat diet (HFD). (A) Timeline of experimental protocol for experiments in Figure 4. Created with Biorender. (B and C) Percentage of body weight change during HFD, before and after stereotaxic surgery injecting AVV2-hsyn-mcherry-cre in DMH, using their body weight before the access to *ad libitum* HFD as baseline (B) Female mice (C) Male mice (B: baseline-corrected followed by two-way ANOVA with multiple comparisons, p>0.9999 n=8 WT mice and 4 DMH KO mice; C: baseline-corrected followed by two-way ANOVA with multiple comparisons, p>0.9999 n=10 WT mice and 5 DMH KO mice). (D and E) Daily HFD cumulative food intake of DIO mice 2 weeks after stereotaxic surgery (D) Female mice (E) Male mice (D: two-way ANOVA with multiple comparisons, p=0.5927 n=8 WT mice and 4 DMH KO mice; E: two-way ANOVA with multiple comparisons, p=0.9409 n=10 WT mice and 5 DMH KO mice). (F and G) Daily percentage of body weight change using saline as the baseline of DIO mice administered with 100mg/kg of semaglutide (F) Female mice (G) Male mice (F: baseline-corrected followed by two-way ANOVA with multiple comparisons, p=0.9882 n=8 WT mice and 4 DMH KO mice; G: baseline-corrected followed by two-way ANOVA with multiple comparisons, p**<0.0090 n=10 WT mice and 5 DMH KO mice). (H and I) Daily HFD cumulative food intake of DIO mice during semaglutide administration (H) Female mice (I) Male mice (H: two-way ANOVA with multiple comparisons, p=0.7029 n=8 WT mice and 4 DMH KO mice; I: two-way ANOVA with multiple comparisons, p*=0.0128, p**=0.0031 n=10 WT mice and 5 DMH KO mice). Data points represent individual mice.

### 2.7 MC3R deletion in dMH enhances weight loss to semaglutide in male diet-induced obese mice

Semaglutide has demonstrated remarkable efficiency in producing weight loss in the context of diet-induced obesity in humans and pre-clinical rodent models(39,40). Therefore, since our previous experiments were performed in lean regular chow fed mice, we next tested if dMH deletion of MC3R contributes to the anorexic response to semaglutide in diet induced obese mice (**Figure 5A**). Male dMH MC3R KO mice displayed an enhanced anorexic and weight loss response following semaglutide administration, relative to WT littermate mice (**Figures 5G and 5I**). However, consistent with our prior studies in lean mice, dMH MC3R deletion in female mice did not alter the anorexic or weight loss response to semaglutide (**Figures 5F and 5H**). Following cessation of semaglutide treatment, the female mice did not show any feeding or body weight differences compared to WT mice (**Figures S7A and S7C**). In contrast, male dMH MC3R-KO mice re-gained weight at a slower pace than WT littermates (**Figure S7B**) and consumed less food in the 24 hours after the cessation of semaglutide treatment (**Figure S7D**). Thus, dMH deletion of MC3R enhances the anorexic response to semaglutide in both lean and obese male mice.

### 2.8 Activation of DMH MC3R neurons acutely alters energy homeostasis

It has been demonstrated that MC3R neurons in the arcuate nucleus exert an orexigenic effect, primarily by regulating the activity of AgRP neurons(44). However, the role of MC3R neurons in medial hypothalamic populations outside of arcuate nucleus has been less well characterized. Given that MC3R deletion in the dorsal portions of the medial hypothalamus (i.e. VMH and DMH) altered energy rheostasis (**Figure 4**), and we observed altered MC3R mRNA expression in the DMH following negative rheostatic challenges (**Figures 2M-2P**), we next utilized chemogenetic activation approaches in MC3R-Cre mice to more precisely characterize the role of DMH and VMH MC3R neurons in energy homeostasis. The chemogenetic activator hM3Dq or control mCherry expressing virus was targeted to MC3R containing neurons in the DMH or the VMH in two separate cohorts of mice. Comprehensive metabolic profiling was performed on these mice following acute stimulation of MC3R neurons in VMH or DMH in both male and female mice (**Figures 6A, 6B, and S8**). Administration of the DREADD agonist CNO (1mg/kg, i.p.) increased cfos expression in hM3Dq expressing neurons, but not mice containing a control mCherry fluorescent protein (**Figures 6C-6F and 6J**), indicating that CNO successfully increases the activity of MC3R neurons. CNO administration to mice expressing hM3Dq in VMH MC3R neurons did not alter locomotion (**Figure 6G**), energy expenditure (**Figure 6H**), or food intake (**Figure 6I**) in male or female mice. Similar null results were obtained in mice expressing control fluorescent protein in VMH MC3R neurons. In contrast to activation of VMH MC3R neurons, activation of DMH MC3R neurons significantly increased locomotion and trended towards increased energy expenditure in female mice (**Figures S9A and S9B**), while in male mice activation of DMH MC3R neurons trended towards increasing locomotion and significantly increased energy expenditure (**Figures S9C and S9D**). When analyzing both sexes combined, activation of DMH MC3R neurons significantly increased locomotion (**Figure 6K**) and energy expenditure (**Figure 6L**) without altering acute food intake (**Figure 6M**).

**Figure 6:**
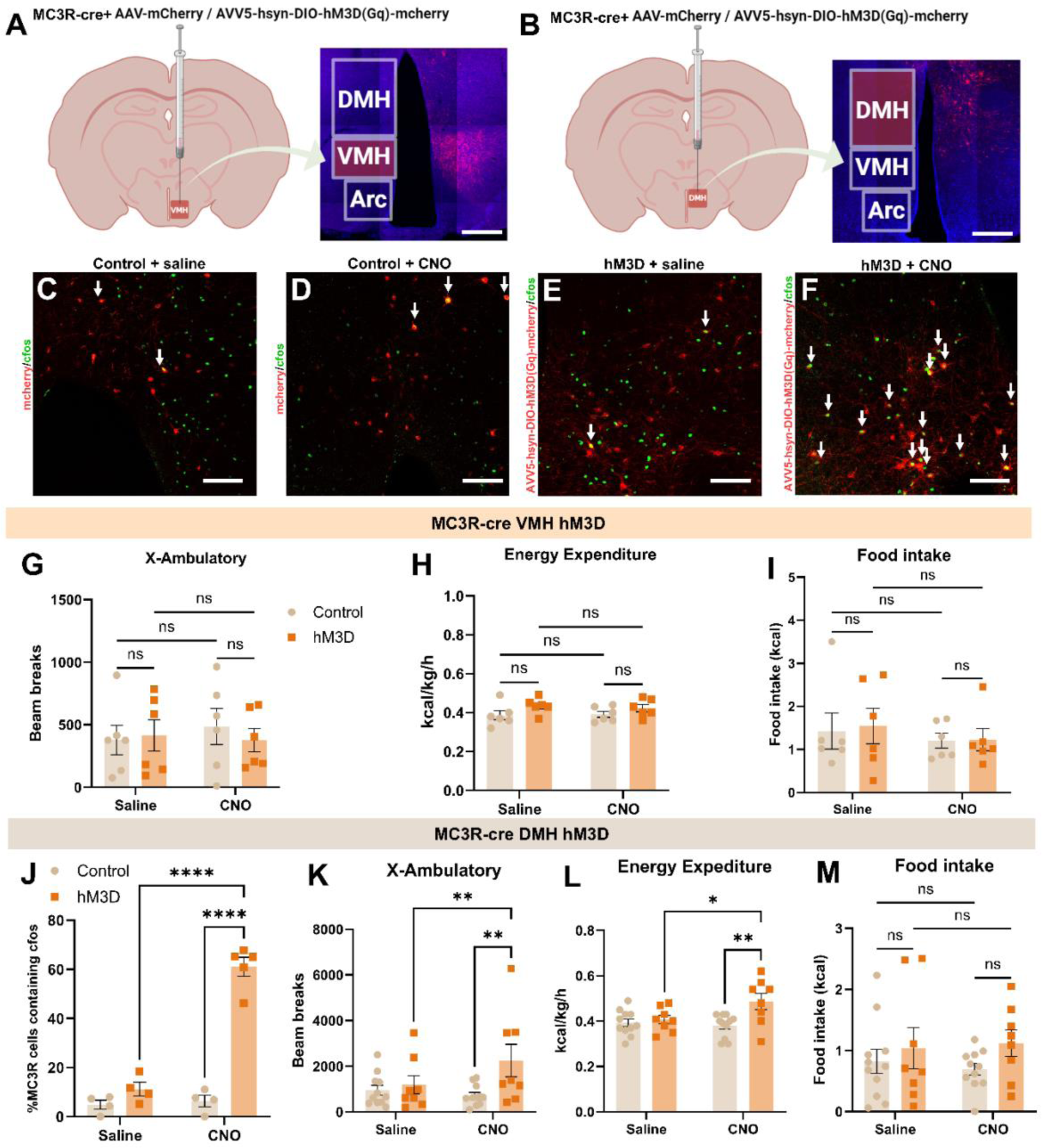
Activation of MC3R expressing cells in DMH (and not VMH) increases energy expenditure and locomotion. (A) Schematics of the stereotaxic surgery location of MC3R-cre mice with unilateral injection of hM3Dq or mcherry viral vector in VMH. Scale bar 300um. Created with Biorender. (B) Schematics of the stereotaxic surgery location of MC3R-cre mice injected with hM3Dq or mcherry viral vector in DMH. Scale bar 300um. Created with Biorender. (C and D) Brain section of a control mouse (DIO-mCherry injected) showing co-localization (white arrows) of cfos (green) and the viral vector (red). Scale bars 100um (C) Administered with saline (D) Administered with CNO. (E and F) Brain section of a hM3Dq mouse that was administered saline showing co - localization (white arrows) of cfos (green) and the viral vector (red). Scale bars 100um. (E) Administered with saline (F) Administered with CNO. (G, H, and I) Control and hM3Dq groups with expression in VMH, after saline or 1mg/kg CNO administration (G) Total locomotion (x-ambulatory) (H) Average of the energy expenditure (I) Total food intake (G: two-way ANOVA with multiple comparisons, p>0.5350, 3 female and 3 male mice/group; two-way ANOVA with multiple comparisons, p>0.2426, 3 female and 3 male mice/group; two-way ANOVA with multiple comparisons, p>0.5079, 3 female and 3 male mice/group). (J) Percentage of MC3R cells that expressed cfos signal, comparing saline and 1mg/kg CNO administrations in both control and hM3Dq groups (two-way ANOVA with multiple comparisons, p****<0.0001). (K, L, and M) Control mCherry and hM3Dq groups with expression in DMH, after saline or 1mg/kg CNO administration (K) Total locomotion (x-ambulatory) (L) Average of the energy expenditure (M) Total food intake (K: two-way ANOVA with multiple comparisons, p**<0.0070, n=5 female/group, 6 control males 3 hM3Dq male mice/group; L: two-way ANOVA with multiple comparisons, p*=0.0142, 5 female and 3 male mice/group; M: two-way ANOVA with multiple comparisons, p**=0.0014, n=5 female/group, 6 control males 3 hM3Dq male mice/group). Data points represent individual mice for all panels.

### 2.9 MC3R cells in DMH are glutamatergic in a sexually dimorphic manner

Although MC3R is expressed in the DMH(38), the specific DMH cell types containing MC3R are unknown. To characterize the DMH cell types expressing MC3R we first attempted to analyze published single cell and single nucleus RNA sequencing datasets of the DMH. However, the low mRNA expression of many receptors critical for energy homeostasis (including MC3R and MC4R) did not allow for adequate detection of MC3R expressing cells (data not shown). Therefore, we instead utilized high sensitivity RNAscope in situ hybridization approaches to map the expression of MC3R containing DMH neurons to previous DMH cell types implicated in energy homeostasis, and to characterize the neurochemical identify of DMH MC3R cells. Prior work has demonstrated a critical role for DMH neurons containing the leptin receptor (LepR) in controlling feeding behavior and circadian rhythms(47,48). To determine if DMH MC3R cells co-localize with DMH LepR neurons, we co-localized mRNA for MC3R and LepR in the DMH of male and female mice. Less than 10% of DMH MC3R neurons co-localized with LepR mRNA, indicating that the vast majority of DMH MC3R neurons do not express the leptin receptor (**Figures 7A-7D**). However, around 30-40% of DMH LepR neurons contain MC3R (**Figure S10A**), suggesting that while the majority of MC3R neurons do not contain LepR, MC3R signaling may still directly regulate the activity of a subset of DMH LepR neurons. Similar co-expression patterns of LepR and MC3R were observed in both male and female mice (**Figure 7D, S10A**).

**Figure 7:**
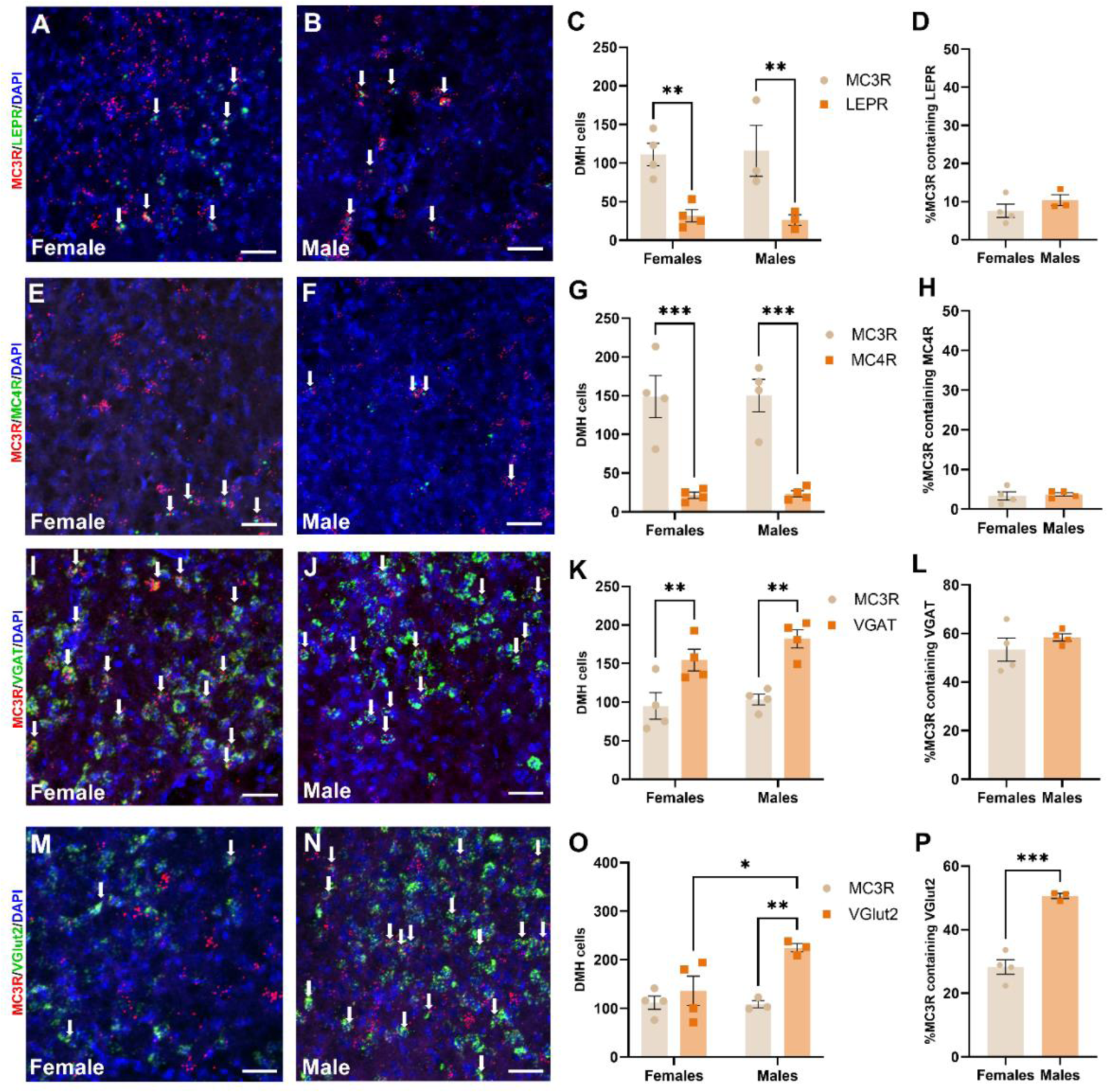
Neurochemical mapping of DMH MC3R neurons in male and female mice. (A and B) Confocal image of the colocalization (white arrows) of MC3R (red) and LEPR (green) mRNA expression with DAPI (blue) in (A) a female mouse brain section (B) a male mouse brain section. Scale bars 40um. (C) Comparison between female and male mice of the number of cells in DMH expressing either MC3R or LEPR mRNA (two-way ANOVA with multiple comparisons, p**<0.0060). (D) Percentage of MC3R cells that co-expressed LEPR of female and male mice (statistical analysis: unpaired t-test, p=0.2894). (E and F) Confocal image of the colocalization (white arrows) of MC3R (red) and MC4R (green) mRNA expression with DAPI (blue) in (E) a female mouse brain section (F) a male mouse brain section. Scale bars 40um. (G) Comparison between female and male mice of the number of cells in DMH expressing either MC3R or MC4R mRNA (two-way ANOVA with multiple comparisons, p***=0.0020). (H) Percentage of MC3R cells that co-expressed MC4R of female and male mice (statistical analysis: unpaired t-test, p=0.7577). (I and J) Confocal image of the colocalization (white arrows) of MC3R (red) and VGAT (green) mRNA expression with DAPI (blue) in (I) a female mouse brain section (J) a male mouse brain section. Scale bars 40um. (K) Comparison between female and male mice of the number of cells in DMH expressing either MC3R or VGAT mRNA (two-way ANOVA with multiple comparisons, p**<0.0071). (L) Percentage of MC3R cells that co-expressed VGAT of female and male mice (statistical analysis: unpaired t-test, p=0.3560). (M and N) Confocal image of the colocalization (white arrows) of MC3R (red) and VGlut2 (green) mRNA expression with DAPI (blue) (M) in a female mouse brain section (N) in a male mouse brain section. Scale bars 40um. (O) Comparison between female and male mice of the number of cells in DMH expressing either MC3R or VGlut2 mRNA (two-way ANOVA with multiple comparisons, p*=0.0111, p**=0.0033). (P) Percentage of MC3R cells that co-expressed VGlut2 of female and male mice (statistical analysis: unpaired t-test, p***=0.0005). Data points represent individual mice for all panels.

Central melanocortin signaling is mediated by both MC3R and MC4R, and DMH MC4R neurons have previously been implicated in energy homeostasis(49,50). Therefore, to quantify the expression of MC3R and MC4R in DMH we performed dual RNAscope *in situ* hybridization for both MC3R and MC4R in male and female mice. MC3R expression vastly outnumbered MC4R expression in DMH at a ratio of nearly 8:1 (**Figures 7E-7G**). Further, less than 4% of DMH MC3R neurons co-express MC4R in both male and female mice (**Figures 7H**). Thus, it is likely that the majority of melanocortin signaling in DMH is mediating by MC3R, although further functional studies are required to confirm this observation.

The DMH consists of a mixture of inhibitory GABA neurons and excitatory glutamate neurons. To determine the neurochemical distribution of DMH MC3R neurons we co-localized MC3R mRNA with either vGAT (GABA marker) or vGLUT2 (glutamate marker). In both males and females, approximately 60% of DMH MC3R neurons co-localized with vGAT (**Figures 7I-7L**). However, while the total number of MC3R positive cells and the total amount of MC3R mRNA transcripts in DMH did not differ between male and female mice (**Figures S10B and S10C**), the percentage of MC3R cells that expressed vGLUT2 was significantly higher in males compared to females (**Figures 7M-7P**). A greater number of vGLUT2 positive cells was observed in the DMH of male vs female animals, suggesting that the increased percentage of MC3R cells containing vGLUT2 in males is driven by an overabundance of vGLUT2 neurons in male mice (**Figure S10D**). Thus, although the total number of MC3R neurons is similar between male and female mice in DMH, the distribution of MC3R in vGAT and vGLUT2 neurons differs between male and female animals.

## 3. Discussion

Emerging evidence indicates that MC3R has a unique function compared to other components of the central melanocortin system (i.e. AgRP, POMC, and MC4R), and other receptors implicated in energy homeostasis (i.e. leptin receptor, glucagon-like 1 receptor, etc.). For example, in contrast to deletion of POMC and MC4R, deletion of MC3R has minimal effects on food intake or body weight in mice(14) or rats(51) provided *ad libitum* access to a standard chow diet. However, MC3R KO mice are hypersensitive to a diverse array of anorexic stimuli including stress-related stimuli (16), anorexic drugs(14,16,17), and tumor-associated anorexia(15,18). Paradoxically, these mice also exhibit an exaggerated response to orexigenic and anabolic stimuli, demonstrating increased weight gain following high fat diet and ovariectomy(14). Together, these findings suggest that MC3R exerts a unique role in energy homeostasis, controlling the magnitude of feeding responses to both orexigenic and anorexic challenges (referred to here as altered energy rheostasis).

MC3R is widely expressed throughout the brain, with expression observed in nearly all major brain regions in mice(38). Of particular importance to energy homeostasis, high MC3R expression is observed in medial hypothalamic regions (arcuate nucleus, ventral-medial hypothalamus, dorsal-medial hypothalamus)(38). Within the arcuate nucleus MC3R is expressed in nearly 100% of AgRP neurons(16), a significant percentage of POMC neurons(16,34,52), and nearly a dozen additional arcuate cell types(16,34). Strong expression is also observed in ventral medial hypothalamus and dorsal medial hypothalamus, although the function and molecular identity of MC3R containing cells in VMH and DMH are less well understood. Here, we show that adult specific deletion of MC3R in the medial hypothalamus (MH) does not change basal feeding or body weight gain when mice are provided *ad libitum* access to regular chow diet (**Figures 1F-1I**). In contrast, like global MC3R KO mice, viral mediated deletion of MC3R in MH recapitulates both the enhanced feeding on HFD (**Figures 1N-1Q**) and the enhanced anorexic response to GLP1R stimulation (**Figures 1J-1M**) observed in global MC3R KO mice. Thus, despite widespread expression of MC3R throughout the brain, many of the core features of energy rheostasis can be recapitulated by selectively deleting MC3R within a small subset of neurons in the medial hypothalamus. Since this study focused on characterizing the role of MH MC3R signaling in the acute response to energy rheostatic stimuli (i.e. semaglutide, HFD, and transition from HFD to regular chow diet), further work is required to characterize the long-term effects of MH deletion of MC3R on the development of diet induced obesity. However, data presented here indicate that reduced/impaired MC3R signaling within the medial hypothalamus alters the magnitude of acute behavioral responses to both orexigenic and anorexic stimuli. Together, these findings support a functional role for medial hypothalamic MC3R signaling in controlling energy rheostasis by regulating the magnitude of feeding responses to both orexigenic and anorexic challenges.

Prior work on the role of MC3R in feeding has primarily focused on the function of MC3R in the arcuate nucleus (14,16), while its role in other medial hypothalamic regions is less well understood. Here, we demonstrate that in addition to the arcuate nucleus, the DMH likely contributes to the role of MC3R in energy rheostasis. For example, deletion of MC3R primarily localized to the DMH enhances the anorexic effects of GLP1R stimulation in male mice (**Figures 4D, 5G, and 5I**) and increases body weight gain and feeding on HFD (**Figures 4F and 4H**). Although MC3R expression was sometimes observed in the nearby VMH, altered energy rheostasis was observed in mice without expression in the VMH (**Figures S4 and S6**), suggesting that VMH may have a less prominent role in energy rheostasis. In some cases, we also observed sparse labeling in the posterior hypothalamus (**Figures S4 and S6**), and we cannot completely exclude a role for this structure in energy rheostasis. Along these lines, further work is required to map the precise sub-regions of DMH (i.e. anterior, medial, and posterior) mediating the role of DMH MC3R signaling in energy rheostasis. However, the functional neuroanatomical experiments presented here indicate that additional medial hypothalamic structures outside of the arcuate nucleus also regulate energy rheostasis.

Consistent with an important role for DMH MC3R signaling in energy homeostasis, mRNA expression of MC3R is reduced in DMH following semaglutide administration (**Figures 2N and 2P**), and selective stimulation of DMH MC3R neurons, but not nearby VMH MC3R neurons, acutely alters energy expenditure and locomotor activity (**Figures 6K and 6L**). Although we did not observe acute effects on feeding following stimulation of DMH MC3R neurons, it is possible that more sustained, long-term activation of DMH MC3R neurons alters feeding behavior. Consistent with this possibility, deletion of MC3R in DMH does not acutely effect feeding in the first 24 hours following behavioral manipulations and requires multiple days following high fat diet or semaglutide administration for effects on feeding to appear (**Figures 4D, 4H, and 5I**). Based on these findings, we conclude that DMH likely also contributes to MC3R-mediated effects on energy rheostasis, primarily by modulating energy expenditure and locomotor activity. We thus propose that MC3R acts via multiplexed neural circuits including the arcuate nucleus and DMH to control energy rheostasis (**Figure S11**), analogous to other receptors involved in energy homeostasis, such as the leptin receptor(53–55) and GLP1R(39,56–58), which mediate their effects by acting on multiple neural pathways. Further work is required to map the activity of DMH MC3R neurons in response to orexigenic and anorexic stimuli and to determine how MC3R signals in DMH neurons.

It is notable that sex differences exist in the energy rheostasis phenotypes reported here in male and female MC3R-KO mice. Both MH and dMH specific deletion of MC3R increases body weight gain following high fat diet in male, but not female mice. Given that female mice are more resistant to high fat diet induced obesity than male animals(59–61), and trends towards increased weight gain were observed in female MC3R KO mice, this difference may be due to inherent sex differences in diet-induced obesity between male and female mice. Conversely, the enhanced weight loss response to semaglutide is observed following MH MC3R deletion in female mice, but not in female mice with dMH deletion of MC3R. Since enhanced anorexia is observed in female global MC3R KO mice(16,44), this suggests that additional MC3R containing regions in MH (i.e. arcuate nucleus or VMH) are required for producing enhanced anorexia in females. Consistent with the sex differences reported here, global MC3R KO mice also exhibit sexually dimorphic feeding and behavioral phenotypes. For example, female MC3R KO mice (but not males) demonstrate an enhanced response to stress-induced anorexia(16) and reduced sucrose preference that is not observed in male MC3R KO mice(62).

In addition to the sexually dimorphic effects of MC3R deletion on behavioral control, the neuroanatomical distribution of MC3R is remarkably sexual dimorphic in mice(38), with increased MC3R expression observed in the arcuate nucleus in male animals, while female mice have higher expression of MC3R in the anteroventral-periventricular nucleus (AVPV), ventral premamillary nucleus (PMv), and the bed nucleus of the stria terminalis (BNST)(38). Although the total amount of MC3R containing cells and overall MC3R expression is the same in the DMH of male and female mice, the distribution of MC3R in glutamatergic, but not GABAergic, neurons is drastically different between the sexes (**Figures 7I-7P and S10D**). Here, we demonstrate that the prevalence of vGLUT2 containing cells is higher in the DMH in male mice (**Figures 7P and S10D**), while the number of vGAT expressing cells in DMH are similar in male and female animals (**Figures 7L and S10D**). As a result, a higher percentage of MC3R containing DMH cells express vGLUT2 in male animals (**Figure 7O**). This neuroanatomical finding offers a putative explanation for the male specific effects on energy rheostasis following dMH deletion of MC3R (**Figures 4 and 5**). Future work is required to precisely map the function of sexually dimorphic DMH MC3R circuits, and to determine how these sex differences contribute to the role of MC3R in energy rheostasis. However, the results presented here support the existence of sexually dimorphic MC3R circuits controlling energy rheostasis and highlight the significance of MC3R circuits outside of the arcuate nucleus in controlling energy rheostasis.

## 4. Methods

### Animals

All animal experiments were approved by the University of Illinois Institutional Animal Care and Use (IACUC) committee. The experiments were performed on littermate mice (6-16 weeks old) that were approximately matched for age and sex between experimental and control groups. MC3R-floxed mice were generated by Taconic using CRISPR-Cas9 gene editing approaches as previously described (13). Heterozygous MC3R-floxed mice were bred together to generate littermate homozygous flox/flox and WT mice that were used for experiments. MC3R-Cre mice were previously described(14). All mice were initially caged in groups of 2 to 5 mice per cage of the same sex until food intake experiments started, when they were single caged for at least 3 days prior to starting experiments. Mice were housed in temperature-controlled (20-21°C) and humidity-controlled cages and the room was under a 12-hour light and 12-hour dark cycle (7 AM-7 PM). All mice had *ad libitum* access to regular chow food and water, unless otherwise noted in the text and figure legends.

### Viral Vectors

For MC3R deletion studies Cre-expressing adeno-associated viral vectors (AAV2-hsyn-mcherry-cre; (UNC GTC Vector #AV6445B) were injected in both WT and MC3R floxed homozygous littermate mice. For chemogenetic activation of DMH and VMH MC3R neurons, the control group (MC3R-Cre positive mice) was unilaterally injected with AAV5-hsyn-DIO-mcherry (Addgene #50459). Experimental mice were injected with AAV5-hsyn-DIO-hM3D(Gq)-mcherry (Addgene #44361).

### Stereotaxic viral injections

Mice were anesthetized in an isoflurane chamber and placed in a stereotaxic frame (kopf) with constant flow of isoflurane. Viral injection coordinates for the MC3R-flox MH-KO injections were: A/P, −1.35mm and −1.8mm (from the bregma); M/L, −0.40mm and +0.40mm; D/V, −5.65mm, −5.75mm, and −5.80mm (from surface of the brain). In each A/P coordinate, 3 injections of viral vectors were delivered in three D/V coordinates, and 75nl, 100nl and 75nl of virus was injected, respectively, in each D/V site at a rate of 50nl/min. Viral injection coordinates for the MC3R-flox dMH-KO and the MC3R-cre chemogentics experiments were: A/P, −1.7mm (from the bregma); M/L, −0.30mm; D/V, −5.10mm (from surface of the brain). The viral vectors were delivered at a rate of 10nl/min, with a total of 60nl (1 side). After the injection, the glass pipette was left in the more dorsal coordinate for 5 additional minutes to prevent leakage of virus from the targeted brain region. Mice were administered 5 mg/kg carprofen subcutaneously after surgery, had their body weights measured, and were returned to their home cages. For all experiments, there was a 2-week period post surgery before starting experiments to allow time for viral expression and recovery from surgery.

### Indirect Calorimetry and locomotion measurements

For chemogenetic experiments, to measure the energy expenditure and x-ambulatory (locomotion), the mice were placed into Comprehensive Laboratory Animal Monitoring System (CLAMS, Columbus instruments, Columbus, US) cages. Mice were habituated to the CLAMS cages for 24 hours before beginning behavioral measurements. Food intake, locomotion, and energy expenditure were automatically calculated every 15 minutes by the CLAMS system. Energy expenditure was calculated as the average per hour and the x-ambulatory as the sum of the values each hour.

### Chemogenetic experiments

All the mice used in this assay were MC3R-cre positive and the stereotaxic viral injections were of either cre-dependent mcherry (control group) or cre-dependent hM3Dq-Cherry viral vectors (experimental group). These experiments were performed in the Comprehensive Laboratory Animal Monitoring System (CLAMS, Columbus instruments, Columbus, US) as described above. On the first day after habituation to the CLAMS, half of the mice were administered saline (200ul, i.p.), and the other half received 1mg/kg CNO (dissolved in 200ul saline, i.p.) one hour before the start of the dark cycle (6pm). On the following day the groups were swapped so that all mice received both saline and CNO. The measurements were analyzed in the first hour of the dark cycle (i.e. 2 hours after the saline or CNO administration).

Validation of the technique was performed by administrating half of the control and experimental groups with saline (0.9% NaCL, 200ul, i.p.) and the other half with CNO (1mg/kg, 200ul, i.p). Perfusion was performed 90 minutes after injections and sectioning and viral location was performed as described in the “*Perfusion, Sectioning, and viral location assessment”* section and followed by “*Immunohistochemistry and cfos quantification*”.

### Perfusion, Sectioning, and viral location assessment

Following experiments all mice were perfused with 10% formalin followed by dissection of the brain. The brains were transferred to a 10% formalin solution for 24 hours, followed by 24 hours in 10%, 20% and 30% sucrose solutions in 1x PBS. Brain slices containing the hypothalamus were then obtained by sectioning on a cryostat (Leica, CM3050S) at 40µm thickness. These sections were then placed into 24 well plates containing 500µL of 1x ultra-pure PBS, mounted on Superfrost glass slides (Fisher), and imaged with confocal microscopy (Zeiss LSM 510 microscope, Z-stack and tile scan of whole hypothalamus). For RNAscope experiments sections were mounted directly onto glass slides (20µm) and RNAscope was performed as described in “*RNAscope in situ hybridization and mRNA quantification”*.

### Immunohistochemistry and cfos quantification

The 40um brain sections were placed into 24 well plates containing 500µL of blocking buffer (100 mL Ultrapure 1X PBS, 2 grams Bovine Serum Albumin, and 100µL of Tween 20) and placed on a shaker at room temperature and allowed to sit for 2 hours. Next, a master mix of the 1:1000 cFos (9F6 Rabbit mAb, Cell Signaling Technology) primary antibody in blocking buffer was prepared. 500µL of this master mix was added to each well-containing brain sections and placed onto a shaker at 4°C overnight. The primary antibody mixture was replaced by 500µL of ultra-pure 1X phosphate-buffered saline (PBS) and placed on a shaker at room temperature for 10 minutes and this step was repeated twice more. The secondary antibodies (Goat anti-Rabbit IgG (H+L) Cross-Adsorbed Secondary Antibody, Alexa Fluor™ 488) were prepared in blocking buffer in a 1:500 concentration and then 500µL of the secondary was added to each well and placed on a shaker, at room temperature, and incubated for 2 hours. Following three 10 -minute wash steps with 1X ultra-pure PBS, the sections were mounted on Superfrost glass slides (Fisher) and analysis of the images were performed using LSM700 confocal microscope (Z-stack, 20x). Duplicate sections with mcherry expression in DMH for each mouse were used to count the mcherry and cfos signals, and the average of those measurements was calculated for each mouse for statistical analyzes. The cfos quantification was performed using ImageJ/Fiji software and the total number of DMH cells expressing MC3R (labeled with mcherry viral vectors) was counted for the entire section and the percentage of these cells co-expressing with cfos was quantified for the statistical analysis.

### RNAscope in situ hybridization and mRNA quantification

RNAscope analysis was performed on WT and MC3R-flox mice, all injected with AVV2-hsyn-mcherry-cre. The animals were perfused as described above, however using 4% paraformaldehyde instead of 10% formalin according to ACD RNAscope recommended protocols, and 20µm sections were directly mounted onto Superfrost glass slides for RNAscope in situ hybridization. RNAscope multiplex fluorescent in situ hybridization version 2 was used according to the protocol described in the kit. MC3R mRNA expression was visualized using the probe Mm-Mc3r-C2 probe (Ref: 412541-C2). Images were obtained via confocal microscopy (Z-stack and tile scan of whole hypothalamus). The mRNA count (**Figures 1C and 1D**) was performed using Fiji/ImageJ software, where a region of interest (ROI) was created and placed on the area with mRNA expression in the MH region. The Mc3r mRNA was manually counted, and the same ROI size was used for all the images.

For MC4R, LEPR, VGAT, VGlut2, AgRP, and POMC the probes utilized were: Mm-Mc4r-C3 (Ref: 319181-C3), Mm-Lepr-C3 (Ref: 402731-C3), Mm-Slc32a1 (Ref: 319191-C1), Mm-Slc17a6 (Ref: 319171-C1), Mm-Agrp-C2 (Ref: 400711-C2), and Mm-Pomc-C3 (Ref: 314081-C3) respectively. The confocal images were taken using LSM700 microscope and Zen software (z-stack and 20x zoom) and the cells were counted in the entire DMH, VMH, or ARC regions (**Figures 2 and S3**) using Fiji/ImageJ software. Duplicate or triplicates sections for the probes combinations were analyzed for each mouse, and the average of those measurements used to establish the number of cells which expressed the mRNA for statistical analysis. The colocalization cell count was performed considering the soma (stained with DAPI) containing at least one transcript of each probe as a positive cell. For the mRNA pixels measurement, using Fiji/Image J, a ROI was created covering entirely DMH, VMH, or ARC sections in duplicates or triplicates for each animal, and the signal was measured using the “measure” software feature. The average of the signal value for each animal was used to determine its signal intensity and used for statistical analysis.

### Feeding behavior assays

#### Body weight and food intake measurements

Two weeks after the stereotaxic surgery, which is the time given for the mice to recover from surgery and for viral expression to occur, we started daily body weight and food intake measurements. For the body weight measurements, the individual mice were measured at the same hour every day. At the same time, a previously measured amount of food was given to the mice and the mice were placed in a new clean cage, to avoid errors with crumbles of food on the bottom of the cage that could interfere with the food intake values. Food intake was measured by weighing the food (either regular chow or high fat diet, depending on the experiment) in the hopper and subtracting from the original food weight.

#### Caloric Restriction studies

We measured the ad libitum food intake and body weight for the mice as described in *<u>Body weight and food intake measurements</u>* prior to the caloric restriction. The average food intake was used as a baseline (4.3g) and the animals were given, for 8 consecutive days, 70% of the amount of regular chow that was normally consumed in 24 hours (3g) and their body weight was measured daily. After the caloric restriction period, the animals received *ad libitum* access to food and their food intake and body weight were measured as described above.

#### Diet-induced obesity (DIO)

WT and MC3R-floxed mice had access to *ad libitum* high fat diet (Research Diet Inc, D12492) for seven weeks prior to surgery to induce obesity. After the DIO period, we performed stereotaxic surgical injections as described in “*Stereotaxic viral injections”* to delete the MC3R in DMH. The mice were continuously provided with high fat diet throughout the entire experiment and food intake was measured as described above.

#### Semaglutide administration

We administered the WT and MC3R-KO mice with saline solution subcutaneously (200ul) for 2 days to acclimate animals to being handled and to the subcutaneous injections. On the days they were given saline, their body weight and food intake were measured as described in *“Body weight and food intake measurements”*. Semaglutide was administered at 100ug/kg daily to the mice subcutaneously for 5 days and body weight and food intake measured daily, as described above.

### Data analysis and Statistics

The animals were regrouped according to the viral location. For the MC3R-flox MH-KO animals, a bilateral viral expression in DMH, VMH and Arc was necessary for the mice to be included in the data analysis. For the MC3R-flox dMH KO mice only mice with bilateral viral expression in the DMH were used. For the chemogenetic assays, the mice had unilateral expression and were regrouped after viral location. The specific viral spread of each animal is further outlined in the supplemental figures (Figures S1, S3, S5, S8, and S10). Specific statistical tests are further outlined in the figure legends. Data was analyzed using Graphpad Prism 10 software and they are shown as Mean ± SEM.

## Data Availability

Original data will be available upon request by contacting the corresponding author.

## Acknowledgement

We thank all members of the Sweeney lab for helpful comments on earlier drafts of the manuscript.

---

**Figure S1:**
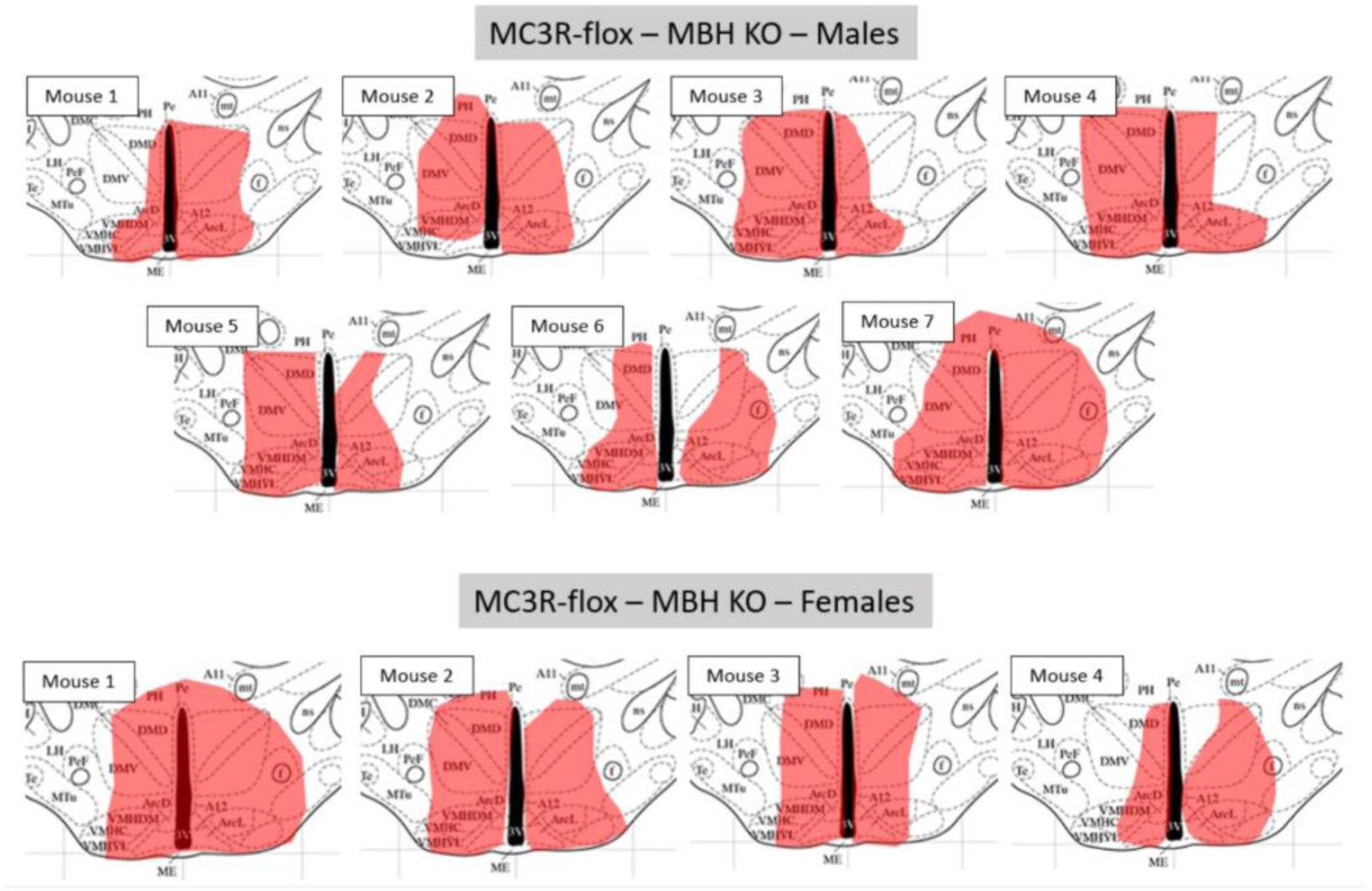
Schematics of the viral expression of all the mice used in the experiments in Figures 1 and S2. Adapted from Allen Brain Atlas - Mouse Brain, using the reference of bregma −1.94mm.

**Figure S2:**
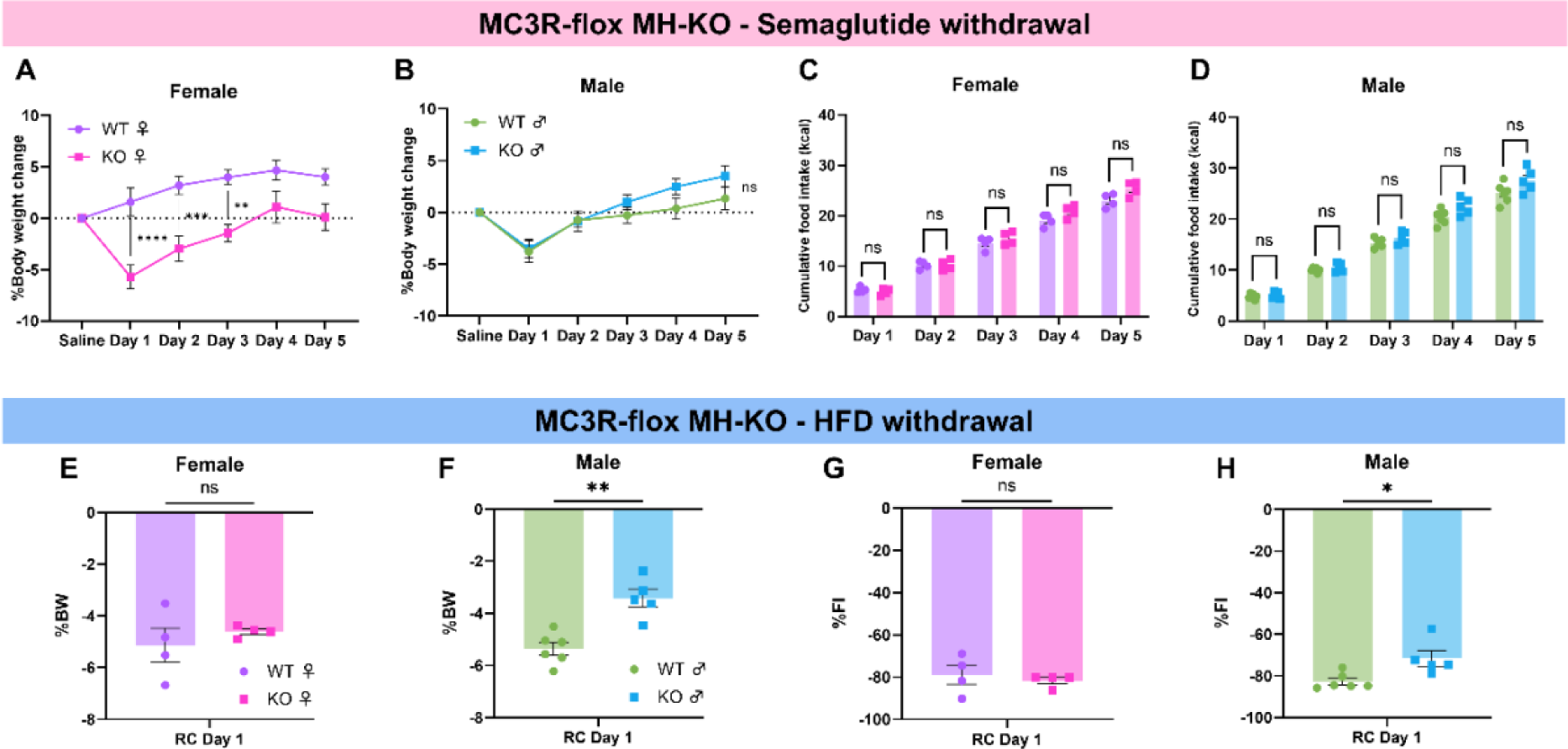
Deletion of MC3R in MH causes female mice to gain significantly less weight after semaglutide withdrawal, while HFD withdrawal causes males to lose less weight. Related to. Figure 1. (A and B) Daily percentage body weight change, using saline as the baseline, during 5 days of semaglutide withdrawal (A) Female mice (B) Male mice (A: baseline-corrected followed by two-way ANOVA with multiple comparisons, p****<0.0001, p***=0.0007, p**=0.0034 n=4 mice/group; B: baseline-corrected followed by two-way ANOVA with multiple comparisons, p=0.3473 n=6 WT mice and 5 MH KO mice). (C and D) 24-hour cumulative food intake during 5 days of semaglutide withdrawal (C) Female mice (D) Male mice (C: two-way ANOVA with multiple comparisons, p=0.2592 n=4 mice/group; D: two-way ANOVA with multiple comparisons, p=0.3738 n=6 WT mice and 5 MH KO mice). (E and F) Daily percentage body weight change, using HFD as the baseline, 24 hours after the change from HFD to regular chow (E) Female mice (F) Male mice (E: baseline-corrected followed by unpaired t-test, p=0.4578 n=4 mice/group; F: baseline-corrected followed by unpaired t-test, p*=0.0010 n=6 WT mice and 5 MH KO mice). (G and H) Daily percentage food intake change, using HFD consumption as the baseline, 24 hours after the change from HFD to regular chow (G) Female mice (H) Male mice (G: baseline-corrected followed by unpaired t-test, p=05871 n=4 mice/group; H: baseline-corrected followed by unpaired t-test, p*=0.0165 n=6 WT mice and 5 MH KO mice). Data points represent individual mice.

**Figure S3:**
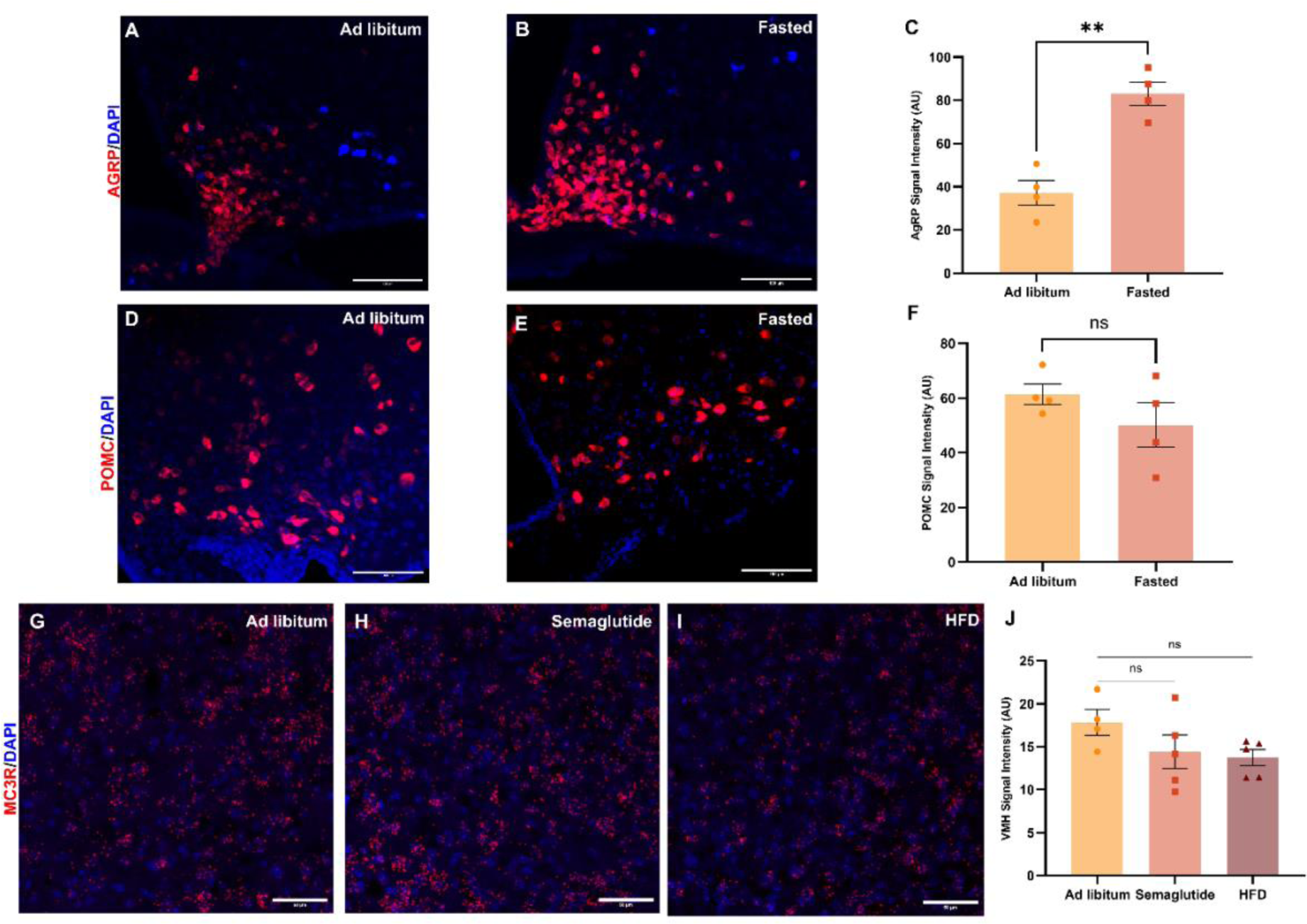
Semaglutide administration and HFD access does not alter MC3R expression in ventromedial hypothalamus (VMH). (A and B) Confocal image of AgRP mRNA expression (red) with DAPI (blue) of a mouse (A) with ad libitum access to regular chow (RC) (B) fasted for 16 hours. Scale bars 100um. (C) Comparison of the intensity of AgRP mRNA signal of mice with Ad libitum access to RC or 16 hours fasted. (unpaired t-test p**=0.0011). (D and E) Confocal image of POMC mRNA expression (red) with DAPI (blue) of a mouse (D) with ad libitum access to RC (E) fasted for 16 hours. Scale Bars 100um. (F) Comparison of the intensity of POMC mRNA signal of mice with Ad libitum access to RC or 16 hours fasted. (unpaired t-test p=0.2587). (G, H, and I) Confocal image of MC3R mRNA (red) expression in VMH with DAPI (blue) of a mouse (G) with ad libitum access to RC (H) after 5 days of semaglutide administration (I) after 5 days of HFD access. (J) Comparison of the intensity of MC3R mRNA signal in ARC during Ad libitum, semaglutide administration for 5 days and access to HFD for 5 days. (unpaired t-test Ad libitum-Semaglutide p=0.2505; Ad libitum-HFD p=0.1564).

**Figure S4:**
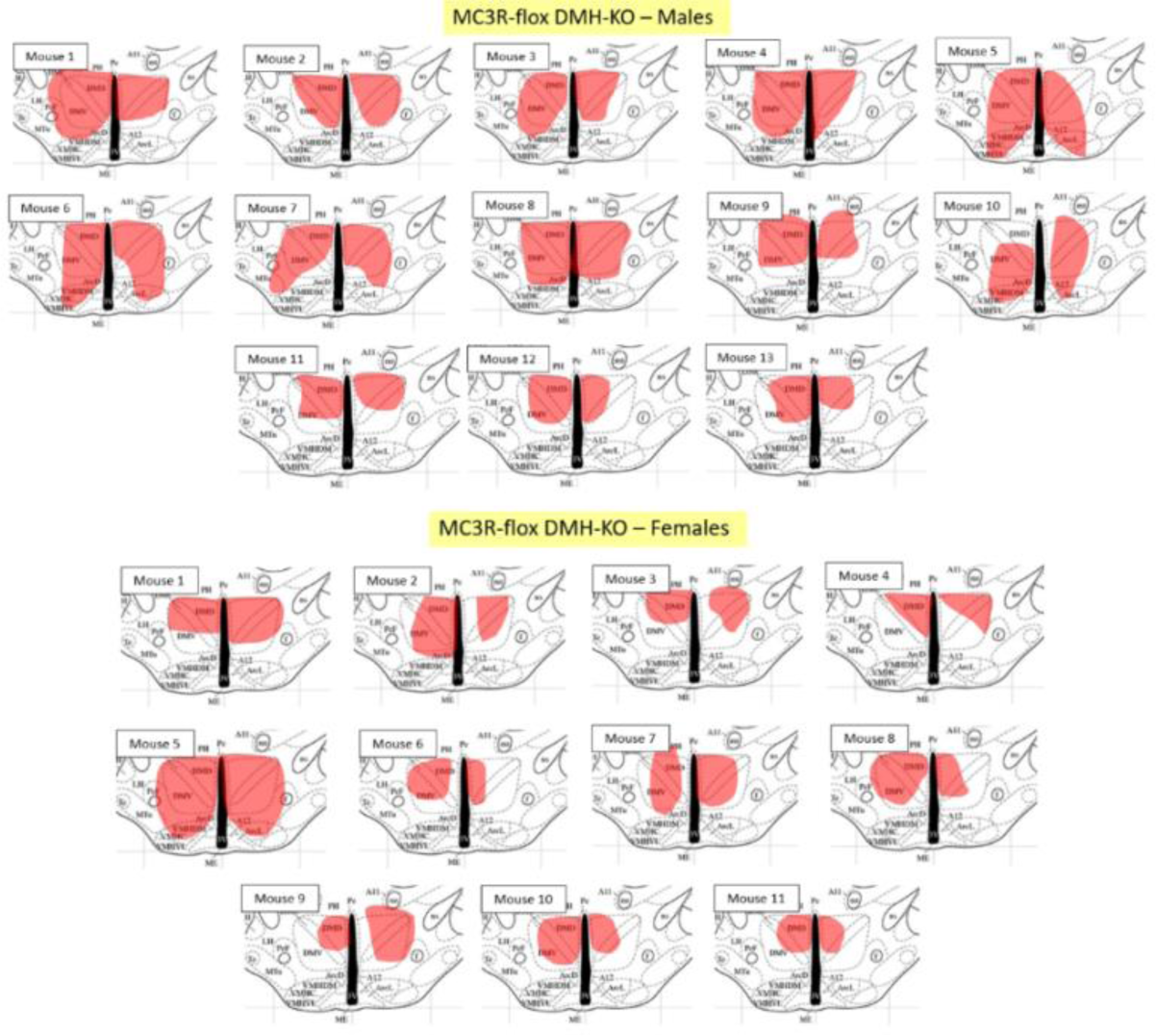
Schematics of the viral expression of all the mice used in the experiments in Figures 2, 3 and S4. Adapted from Allen Brain Atlas - Mouse Brain, using the reference of bregma −1.94mm.

**Figure S5:**
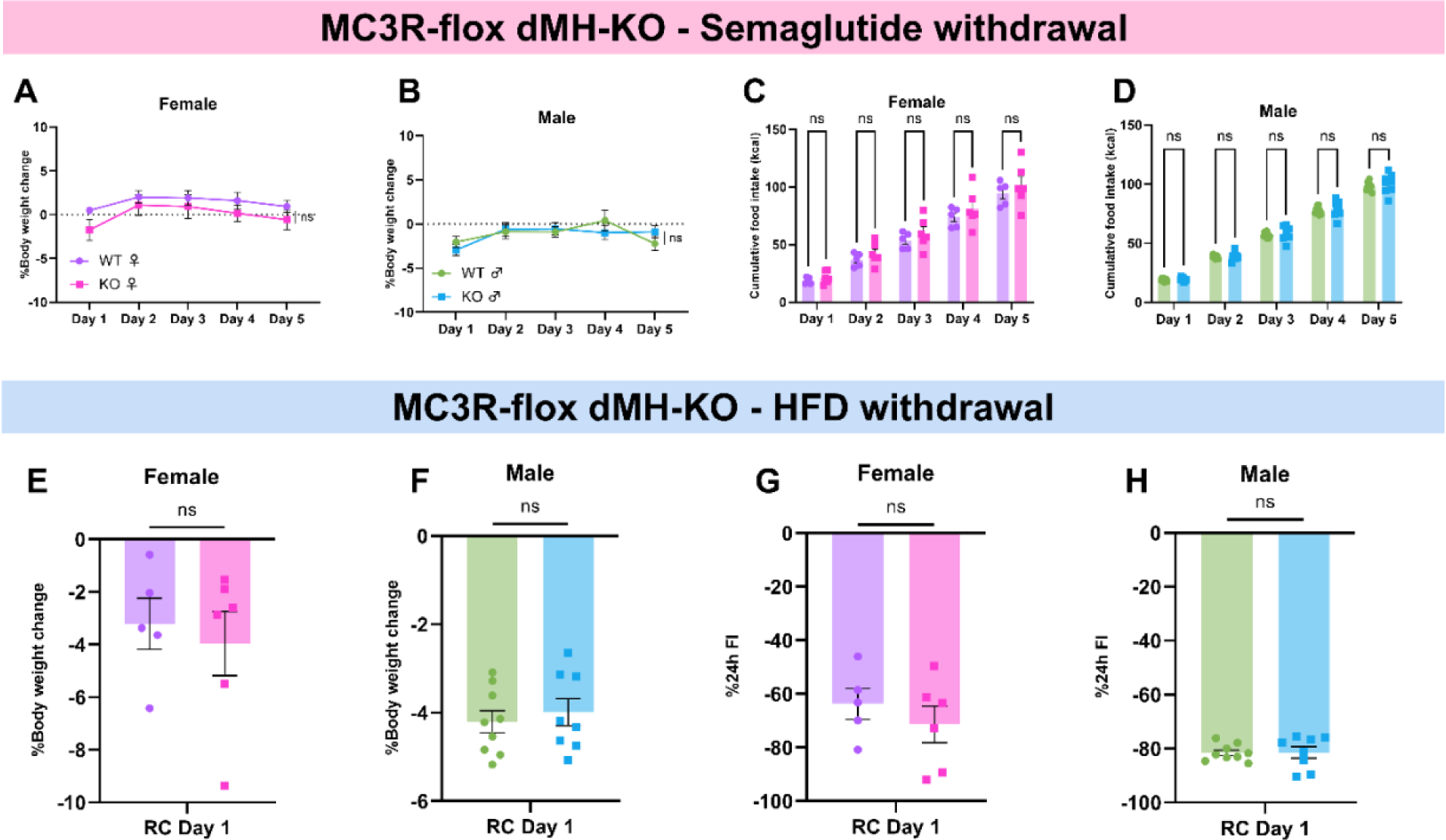
Deletion of MC3R in DMH does not influence weight re-gain or food intake during semaglutide or HFD withdrawal. Related to Figures 2 and 3. (A and B) Daily percentage body weight change, using saline as the baseline, during 5 days of semaglutide withdrawal (A) Female mice (B) Male mice (A: baseline-corrected followed by two-way ANOVA with multiple comparisons, p=0.8293 n=5 mice in WT group and 6 mice in KO group; B: baseline-corrected followed by two-way ANOVA with multiple comparisons, p=0.7163 n=9 WT mice and 8 dMH KO mice). (C and D) 24-hour cumulative food intake during 5 days of semaglutide withdrawal (C) Female mice (D) Male mice (C: two-way ANOVA with multiple comparisons, p=0.9523 n=5 mice in WT group and 6 mice in KO group; D: two-way ANOVA with multiple comparisons, p=0.9660 n=9 WT mice and 8 dMH KO mice). (E and F) Daily percentage body weight change, using HFD as the baseline, 24 hours after the change from HFD to regular chow (E) Female mice (F) Male mice (E: baseline-corrected followed by unpaired t-test, p=0.6510 n=5 mice in WT group and 6 mice in KO group; F: baseline-corrected followed by unpaired t-test, p=0.5933 n=9 WT mice and 8 dMH KO mice). (G and H) Daily percentage food intake change, using HFD consumption as the baseline, 24 hours after the change from HFD to regular chow (G) Female mice (H) Male mice (G: baseline-corrected followed by unpaired t-test, p=0.4221 n=5 mice in WT group and 6 mice in KO group; H: baseline-corrected followed by unpaired t-test, p=0.9405 n=9 WT mice and 8 dMH KO mice). Data points represent individual mice.

**Figure S6:**
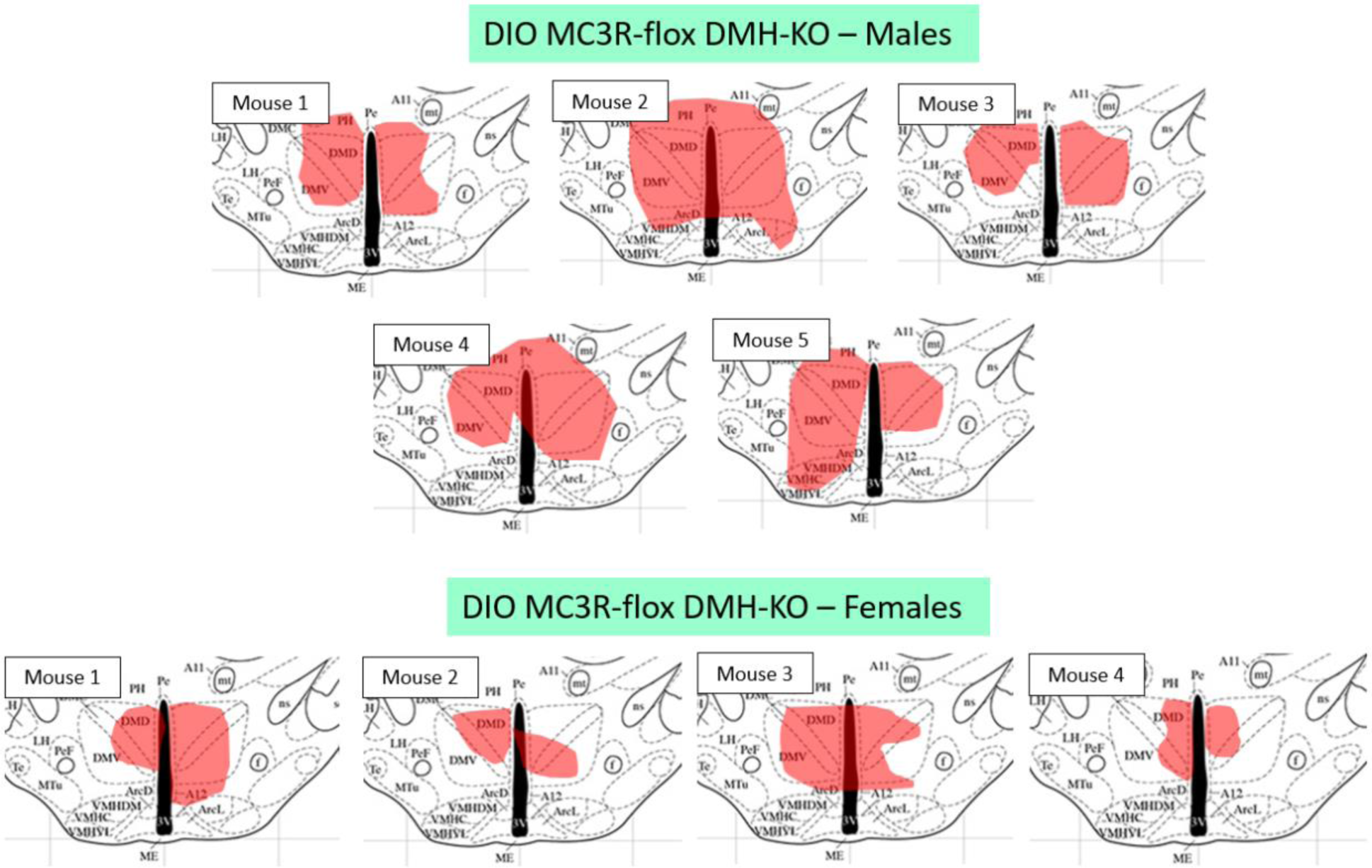
Schematics of the viral expression of all the mice used in the experiments in Figures 4 and S6. Adapted from Allen Brain Atlas - Mouse Brain, using the reference of bregma −1.94mm.

**Figure S7:**
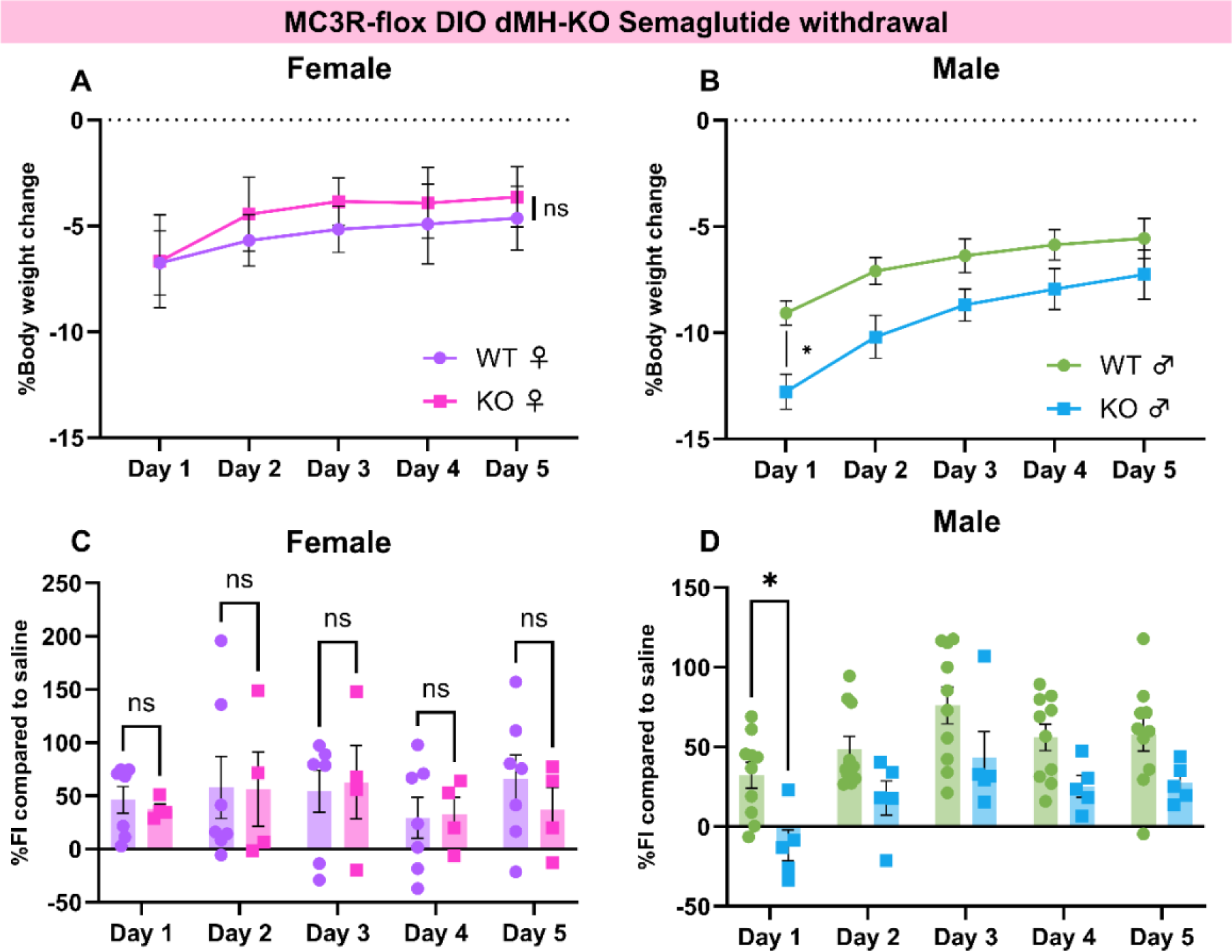
Deleting MC3R in DMH of DIO mice impairs their weight re-gain and food intake in the first day of semaglutide withdrawal. Related to Figure 5. (A and B) Daily percentage body weight change, using saline as the baseline, during 5 days of semaglutide withdrawal (A) Female mice (B) Male mice (A: baseline-corrected followed by two-way ANOVA with multiple comparisons, p=0.9048 n=7 WT mice and 4 dMH DIO KO mice; B: baseline-corrected followed by two-way ANOVA with multiple comparisons, p*=0.0217 n=10 WT mice and 5 dMH DIO KO mice). (C and D) 24-hour percentage food intake difference, compared to saline, during 5 days of semaglutide withdrawal (C) Female mice (D) Male mice (C: baseline-corrected followed by two-way ANOVA with multiple comparisons, p=0.5003 n=7 WT mice and 4 dMH DIO KO mice; D: baseline-corrected followed by two-way ANOVA with multiple comparisons, p=0.0270 n=10 WT mice and 5 dMH DIO KO mice). Data points represent individual mice.

**Figure S8:**
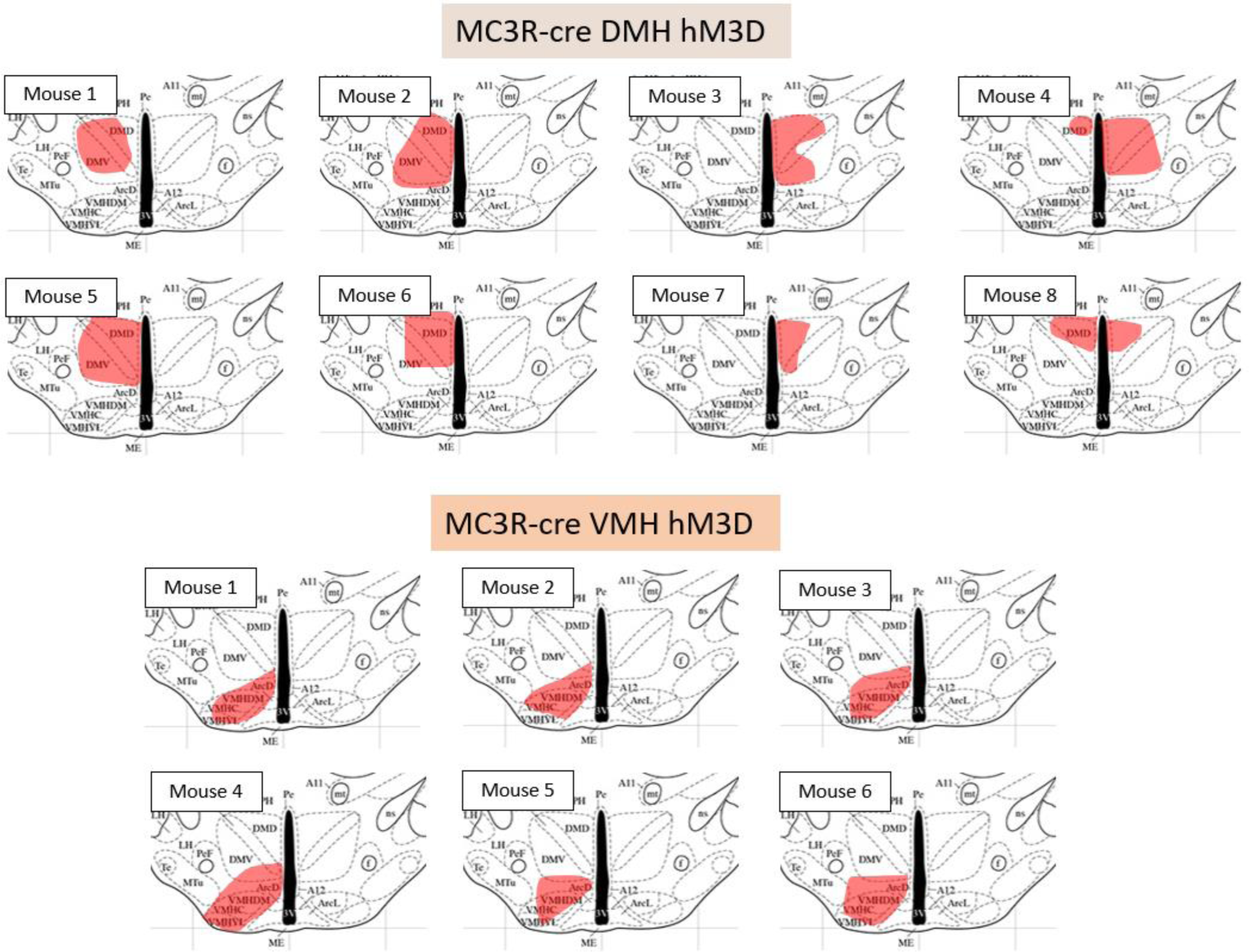
Schematics of the viral expression of all the mice used in the experiments in Figures 6 and S9. Adapted from Allen Brain Atlas - Mouse Brain, using the reference of bregma-1.94mm.

**Figure S9:**
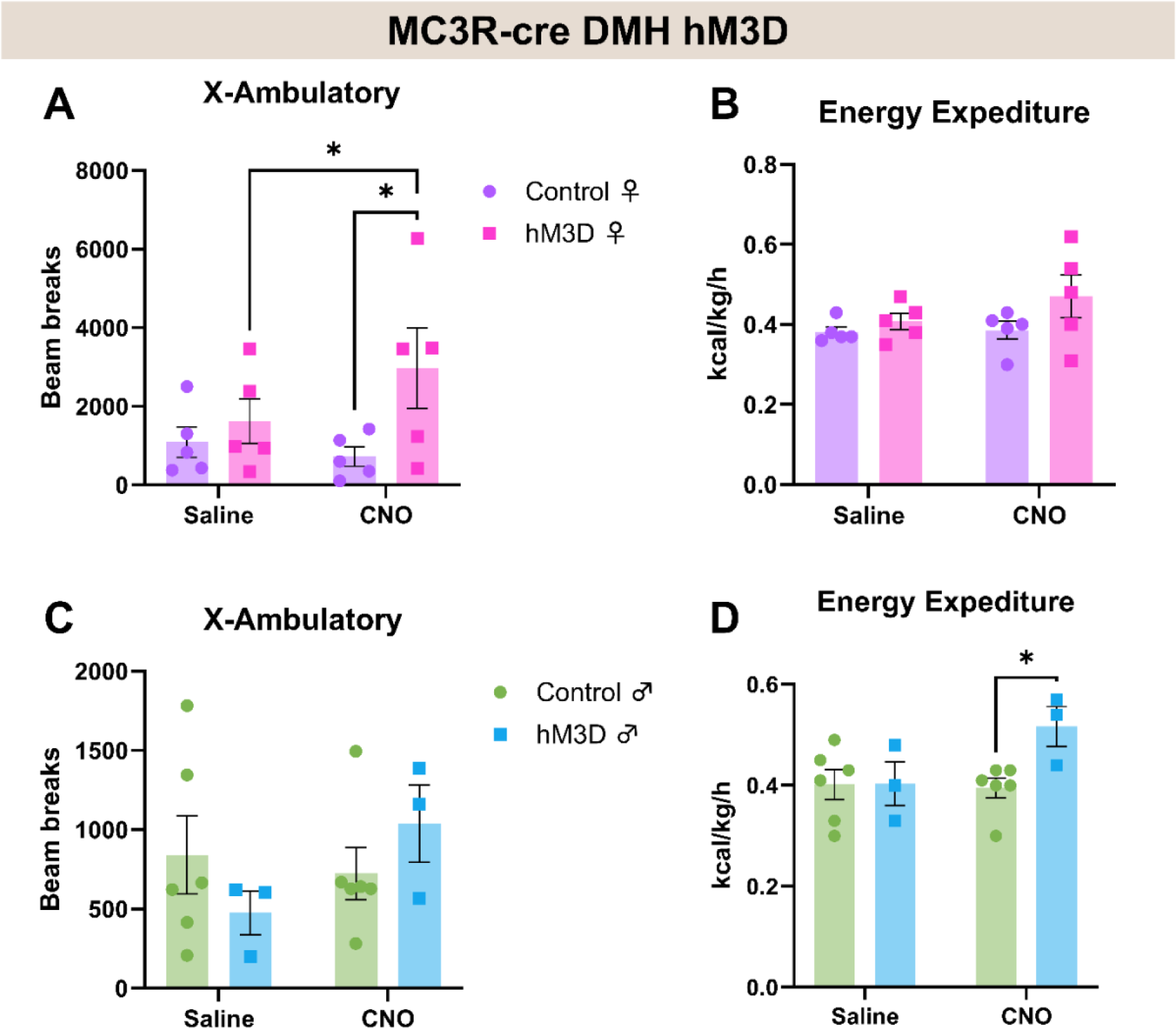
Activation of MC3R cells in DMH significantly increases locomotion in female mice and energy expenditure in male mice. Related to figure 6. (A and C) Total 1 hour locomotion (x-ambulatory) of control and hM3Dq groups with expression in DMH, after saline or 1mg/kg CNO administration (A) Female mice (C) Male mice (A: two-way ANOVA with multiple comparisons, p*<0.0223; C: two-way ANOVA with multiple comparisons, between saline and CNO for the hM3Dq group p=0.0908 n=5/group; between control and hM3Dq receiving CNO p=0.0781 n=6 control and 3 hM3Dq). (B and D) Total 1 hour energy expenditure in control and hM3Dq groups with expression in DMH, after saline or 1mg/kg CNO administration (B) Female mice (D) Male mice (B: two-way ANOVA with multiple comparisons, between saline and CNO for the hM3Dq group p=0.0697; between control and hM3Dq receiving CNO p=0.3592 n=5/group; D: two-way ANOVA with multiple comparisons, p*=0.0188 n=6 control and 3 hM3Dq). Data points represent individual mice.

**Figure S10:**
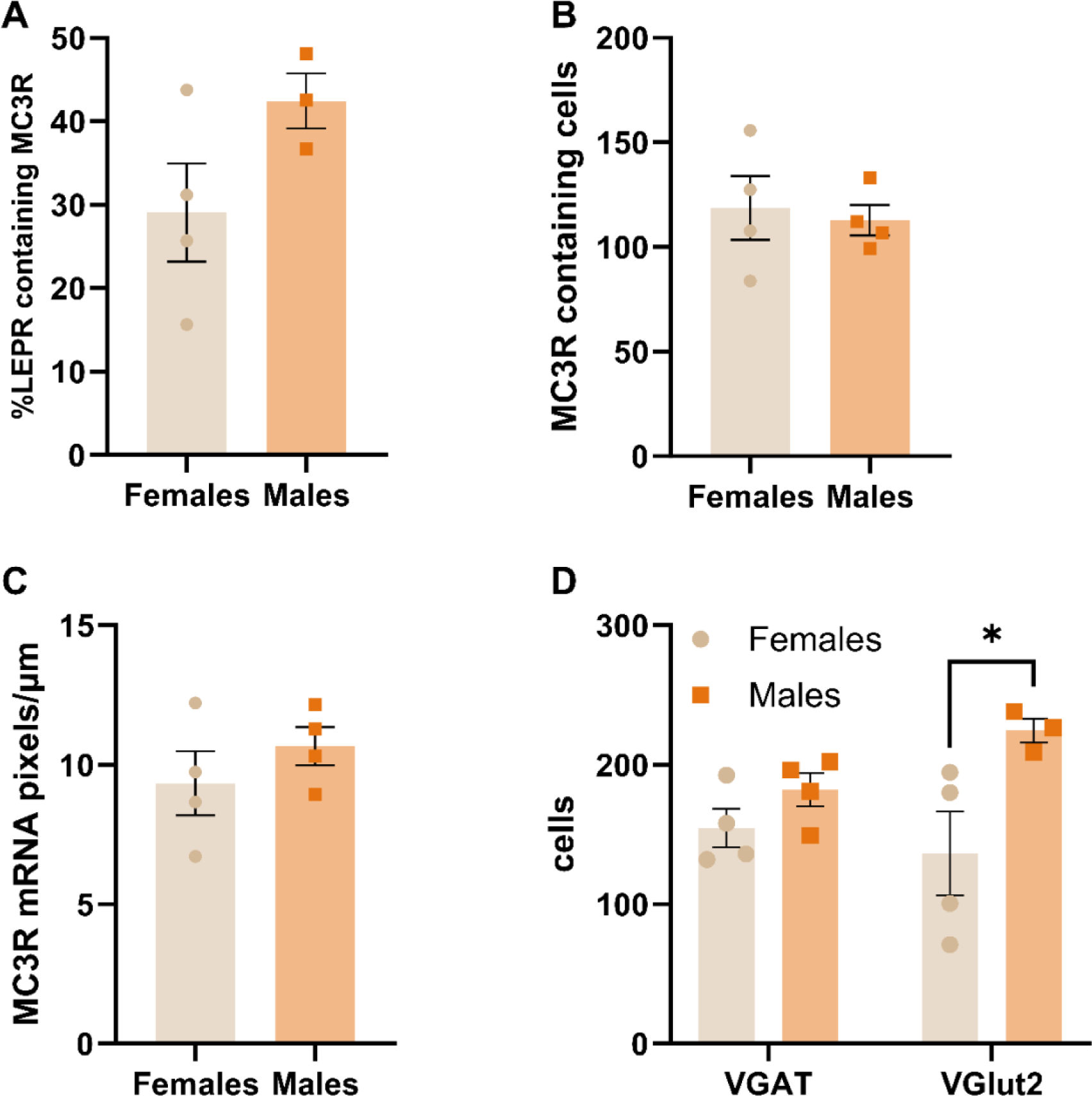
Characterization of DMH MC3R neurons. (A) Percentage of LEPR cells that co-expressed MC3R of female and male mice (statistical analysis: unpaired t-test, p=0.1329). (B) Measurement of the number of MC3R containing cells in males and females in DMH. (C) Density of MC3R mRNA pixels in male and female mice in DMH (statistical analysis: unpaired t-test, p=0.7412). (D) Total number of cells in female and male DMH expressing VGAT or VGlut2 mRNA (statistical analysis: two-way ANOVA with multiple comparisons, p*=0.0102). Data points represent individual mice.

**Figure S11:**
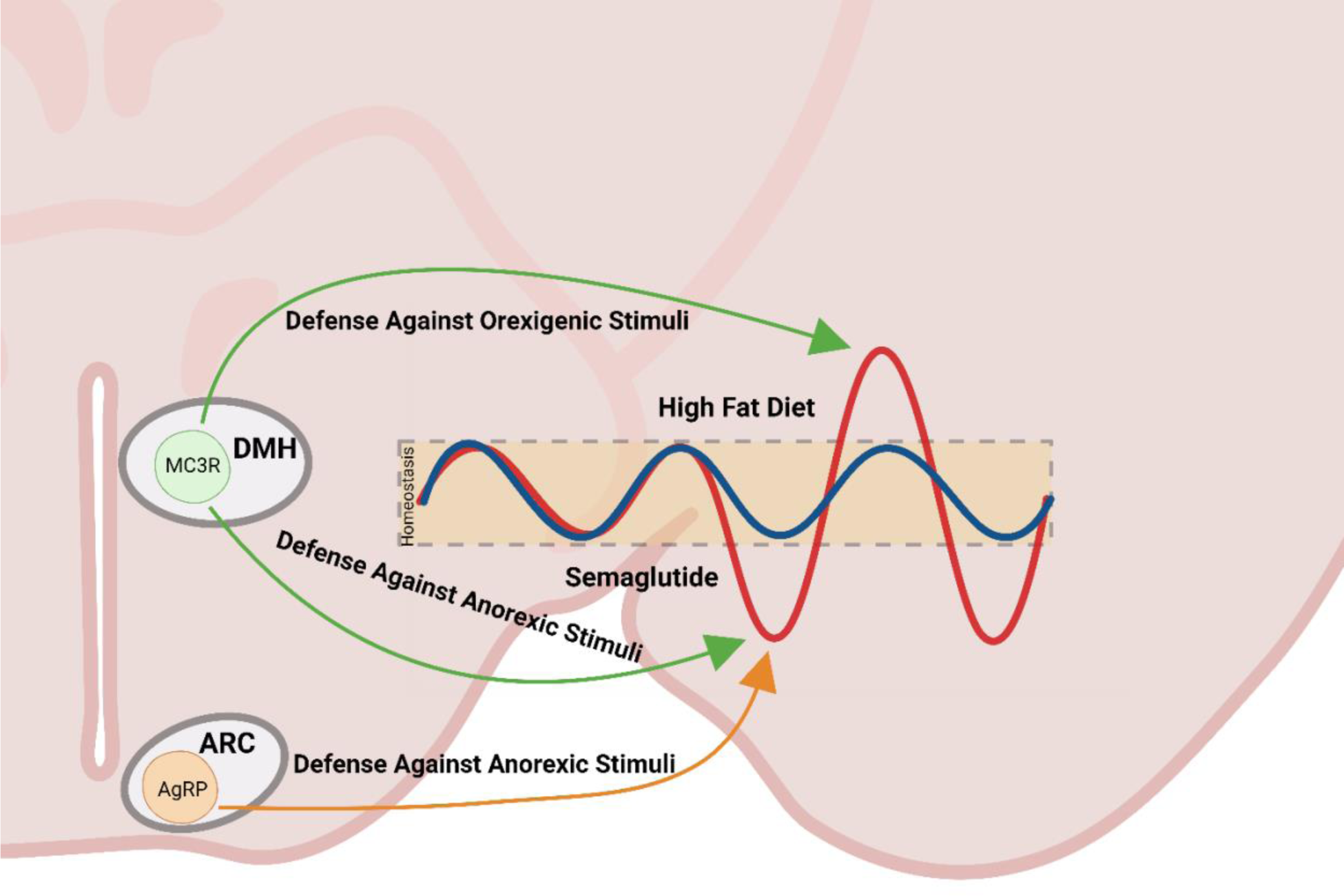
Model of MC3R circuits regulating energy rheostasis. Created with Biorender.

